# Huntingtin-KIF1A-mediated axonal transport of synaptic vesicle precursors influences synaptic transmission and motor skill learning in mice

**DOI:** 10.1101/2022.08.14.503885

**Authors:** Hélène Vitet, Julie Bruyère, Hao Xu, Jacques Brocard, Yah Sé Abada, Benoît Delatour, Chiara Scaramuzzino, Laurent Venance, Frédéric Saudou

## Abstract

Neurotransmitters are released at synapses by synaptic vesicles (SVs), which originate from SV precursors (SVPs) that have traveled along the axon. Because each synapse maintains a pool of SVs, only a small fraction of which are released, it is unclear whether axonal transport of SVPs modifies synaptic function. Here, studying the corticostriatal network both in microfluidic devices and in mice, we find that phosphorylation of the Huntingtin protein (HTT) causes it to recruit the kinesin motor KIF1A, which in turn increases axonal transport of SVPs and synaptic glutamate release. In mice, constitutive HTT phosphorylation leads to SV over-accumulation at synapses, increases the probability of SV release, and impairs motor skill learning on the rotating rod. Silencing KIF1A in these mice restored SV transport and motor skill learning to wild-type levels. Axonal SVP transport within the corticostriatal network thus influences synaptic plasticity and motor skill learning.

## Introduction

Synaptic plasticity underlies our ability to learn. The number of synaptic vesicles (SVs) and release sites at synapses, the probability of the SVs releasing neurotransmitter, and the SV quantal size all affect synaptic strength and thus memory (Katz, 1969). SVs actually begin life as SV precursors (SVPs), which are formed in the cell body and transported along the axon to the presynapse—a distance that can span meters (Guedes-Dias and Holzbaur, 2019; Rizalar et al., 2021; Rizzoli, 2014). It seems intuitively obvious that this long-distance axonal transport influences SV homeostasis, because synapses must be replenished somehow with new vesicles as they release neurotransmitters. Yet the synaptic SV pools contain an average of 400 to 500 vesicles, of which only a few percent participate in synaptic release (Reshetniak and Rizzoli, 2021); these synaptic pools almost certainly serve to ensure that there are always sufficient SVs ready to be released, even with prolonged neuronal stimulation (Denker et al., 2011; Rizzoli, 2014). Moreover, neighboring synapses appear to share localized pools of SVs that circulate between them (Wong et al., 2012).

The idea that SVP transport affects synaptic neurotransmitter release derives from studies of mutations that strongly affect SVP transport, neuronal transmission, and behavior in mice, flies, and worms. In *Caenorhabditis elegans*, null mutants for the kinesin-related gene *unc-104* or the vesicle-associated protein SAM-4 lead to defects in SVP transport, with a consequent lack of SV at synapses and locomotor deficits (Hall and Hedgecock, 1991; Zheng et al., 2014). In *Drosophila*, deletion of the *imac* gene, a kinesin-3 family member, impairs SVP axonal transport and the formation of synaptic boutons (Pack-Chung et al., 2007). In mice, loss of function of *unc-104*’s mammalian homologue, KIF1A (Okada et al., 1995), leads to the accumulation of SVPs in the cell body and a dramatic reduction in the number of SVs at synapses, along with sensorimotor deficits and early postnatal death (Yonekawa et al., 1998). The consequences of completely blocking a molecular motor, however, may not tell us whether enhancement or attenuation of axonal transport influences synapse homeostasis, synaptic transmission, or the function of specific brain circuits. In fact, studies of mice with either constitutively phosphorylated or unphosphorylatable Huntingtin (HTT), a protein that plays a prominent role in axonal transport (Saudou and Humbert, 2016), did not reveal any obvious behavioral phenotypes despite affecting axonal transport (Thion et al., 2015). To what extent, then, does axonal transport of SVPs influence SV release, synaptic strength, and synaptic homeostasis?

We decided to address this question through the Huntingtin (HTT) protein and by revisiting behavior in the mouse models bearing different versions of the serine residue (S421) that needs to be phosphorylated in order for HTT to promote transport toward the synapse. HTT scaffolds various cargoes—endosomes, autophagosomes, vesicles containing BDNF, APP, etc. (Bruyere et al., 2020; Cason et al., 2021; Caviston et al., 2011; Fu and Holzbaur, 2014; Gauthier et al., 2004; Her and Goldstein, 2008; Liot et al., 2013; Wong and Holzbaur, 2014)—along with the appropriate molecular motors for anterograde or retrograde transport and any adaptor proteins that may be needed. HTT contains a polyglutamine tract that, when expanded beyond a certain length, leads to Huntington disease (HD), in part through disruptions in axonal transport (Saudou and Humbert, 2016). In both flies and mice, mutant HTT or its N-terminal fragments induce the formation of axonal aggregates that potentially affect homeostasis of molecular motors, thus disrupting trafficking of SVs (Gunawardena et al., 2003; Li et al., 2003; Weiss and Littleton, 2016). Whether wild-type mammalian HTT regulates the trafficking of SVPs, however, is unclear.

Because the direction of HTT-mediated transport is dictated by S421 phosphorylation (Bruyere et al., 2020; Colin et al., 2008; Ehinger et al., 2020; Vitet et al., 2020), we investigated how phosphorylation of HTT affects axonal transport of SVPs using a reconstituted neuronal circuit on-a-chip. We then studied the effects of S421 mutants that are either constitutively phosphorylated or unphosphorylatable on physiology and behavior in mice and find that phosphorylated HTT recruits the molecular motor KIF1A to promote SVP transport. Our data reveal a functional link between anterograde transport of SVPs within corticostriatal projecting neurons, the synaptic SV pools, and the release probability of SVs at corticostriatal synapses, with consequences for motor skill learning.

## Results

### Huntingtin constitutive phosphorylation increases SVPs anterograde transport and synaptic glutamate release

To investigate whether HTT and its phosphorylation affect the transport of SVP in a physiologically relevant system, we reconstituted mature corticostriatal circuits in microfluidic devices (Moutaux et al., 2018; Virlogeux et al., 2018). These devices have been used to establish HTT’s role in the transport of organelles such as BDNF-containing vesicles and signaling endosomes (Gauthier et al., 2004; Liot et al., 2013; Scaramuzzino et al., 2022; Virlogeux et al., 2018). Microfluidics consist of a presynaptic and a postsynaptic compartment containing cortical and striatal neurons, respectively, and a middle synaptic compartment that receives axons from cortical neurons and dendrites originating from striatal neurons (**Fig. 1A**). The three compartments are connected by 3-μm-wide microchannels that are 500 μm long for axons and 75 μm long for dendrites (Lenoir et al., 2021). The number of striatal axons reaching the synaptic chamber at maturity is limited by the generation of a laminin gradient from the cortical chamber to the striatal chamber, whilst the poly-D-lysine concentration is kept constant (Scaramuzzino et al., 2022). In this configuration, isolated cortical axons unilaterally connect postsynaptic striatal dendrites in the middle compartment enriched in functional synaptic contacts (Ehinger et al., 2020; Moutaux et al., 2018; Scaramuzzino et al., 2022; Virlogeux et al., 2018).

**Figure 1.**
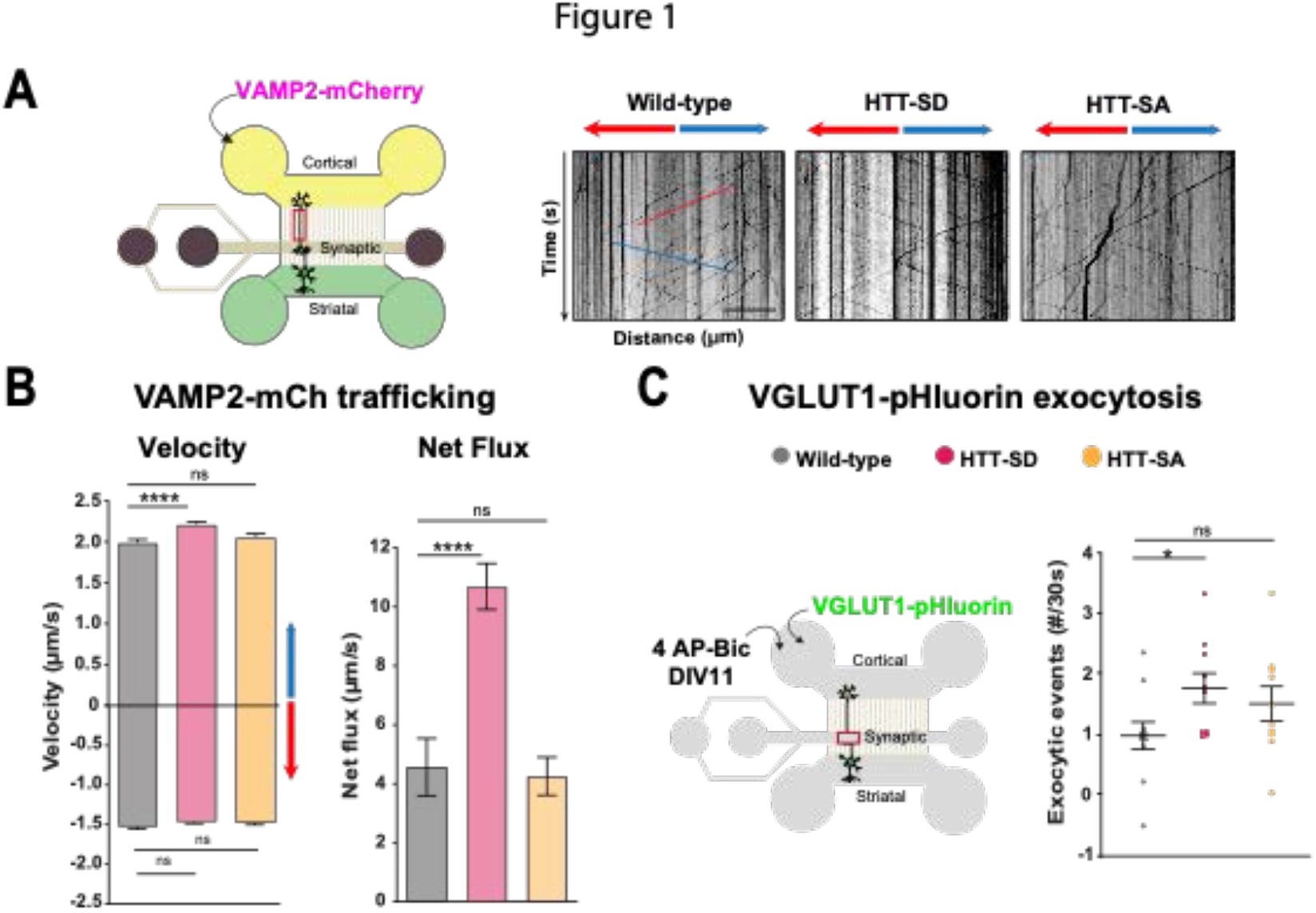
HTT phosphorylation at S421 increases SVP anterograde axonal transport and SV exocytosis. (**A**) Schematic of the 3-compartment microfluidic device allowing the reconstitution of a corticostriatal mature network compatible with live-cell imaging of axons. Cortical axons grow in the cortical chamber (yellow) and connect with the striatal dendrites in the striatal chamber (green) through synapses in the synaptic compartment (purple). On the right, representative kymographs of VAMP2-mCherry vesicle transport in axons for each genotype. Scale bar = 25 µm. (**B**) Segmental anterograde (anterograde: **** p < 0.0001, N=WT:1078 vesicles, HTT-SD: 1886 vesicles and HTT-SA: 1384 vesicles), retrograde velocities (n.s= non significant; WT: 1029 vesicles, HTT-SD: 1564 vesicles, HTT-SA: 2019 vesicles) and directional net flux (**** p<0.0001; N=WT:118 axons, HTT-SD: 157 axons, HTT-SA: 132 axons) of VAMP2-mCherry vesicles. Histograms represent means +/-SEM of 3 independent experiments. Significance was determined using one-way ANOVA followed by a Dunn’s multiple comparison test. (**C)** Schematic of the 3-compartment microfluidic device. Cortical neurons were infected with a lentivirus expressing VGLUT1 together with a pH-sensitive variant of GFP (pHluorin). Stimulation was done with 4 AP bicuculline at DIV11. The number of VGLUT-1 pHluorin exocytosis events within the synaptic chamber of the corticostriatal network, as compared to that of WT and to that of non-stimulated condition (*p<0.05; 6712 events in N=WT, 4640 events in HTT-SD and 5176 events in HTT-SA neuronsHistograms represent means +/-SEM of 3 independent experiments using a total of 11 WT, 10 HTT-SD and 10 HTT-SA seeded microfluidics devices with at least 3 fields per microfluidics device. Significance was determined using a t-test.

We generated cortical and striatal neurons from embryonic day 15.5 (E15.5) wild-type (WT) mice and mice bearing either constitutively phosphorylated (HTT-SD mice) or unphosphorylatable HTT (HTT-SA mice). HTT phosphorylation at serine 421 is mimicked by the replacement of the serine by an aspartic acid, which maintains the positive charge (S421D), whereas the unphosphorylatable form of HTT is obtained by mutating the serine into an alanine (S421A) (Thion et al., 2015). We next transduced cortical presynaptic neurons at a day *in vitro* 0 (DIV0) with a lentivirus encoding the major SNARE protein of synaptic vesicles (v-SNARE), VAMP2 fused to mCherry protein (VAMP2-mCherry), a member of the vesicle-associated membrane protein (VAMP)/synaptobrevin family which labels SVPs (Pennuto et al., 2003). By DIV 10-12, the circuit achieves functional maturity as defined by the kinetics of neurite outgrowth, synapse formation, neuronal transport and synchronous activity, allowing the establishment of functional excitatory connections transmitting information from cortical to striatal neurons (Moutaux et al., 2018). We therefore performed all experiments in the microfluidic devices at this time point. We used high-resolution spinning confocal videomicroscopy to record VAMP2-mCherry particles in the distal part of cortical axons (**Fig. 1A**) and generated kymographs to trace the movement of vesicles (**Fig. 1A**, right panels).

HTT phosphorylation (S421D) increased the anterograde velocity of VAMP-positive vesicles (**Fig. 1B**, left graph), the number of anterograde vesicles, and the linear flow rate (**Supplementary Fig.1A**), leading to an increase in the net directional flux of VAMP2-mCherry vesicles traveling towards the presynapse (**Fig. 1B**, right graph). There was no significant effect on VAMP2-mCherry vesicle velocities or net directional flux of HTT-SA mutation in axons (**Fig. 1B**). HTT phosphorylation at S421 therefore influences transport of SVPs towards the presynapse.

We next investigated whether increased presynaptic anterograde transport of SVPs affects the capacity of presynaptic neurons to release glutamate from SVs at the corticostriatal synapses. We transduced presynaptic cortical neurons with a lentivirus encoding the indicator of vesicle release and recycling VGLUT1-pHluorin, thanks to the fusion of a pH-sensitive GFP (pHluorin) to the vesicular glutamate transport VGLUT1 (Fernandez-Alfonso and Ryan, 2008). We then treated the presynaptic compartment with 4AP/bicuculline at DIV 10-12 to induce neuronal activity and measured the number of exocytic events per active synapse by recording fluorescence within the synaptic compartment (**Fig. 1C**). The amplitude of VGLUT1 events was similar in WT, HTT-SA, and HTT-SD neurons (**Supplementary Fig.1B**), but phosphorylated HTT significantly increased the frequency of release events at synapses (**Fig. 1C**). HTT phosphorylation thus promotes axonal transport of SVPs and increases the capacity of synapses to release glutamate.

### HTT constitutive phosphorylation at S421 impairs motor skill learning in mice

Because we had previously found no differences between HTT-SA, HTT-SD, and WT mice in motor coordination, forelimb strength (grip test), or anxious-depressive behavior (in the elevated plus maze test) (Ehinger et al., 2020), we decided to reassess the three genotypes on the rotarod with a focus on longer-term motor learning rather than mere coordination (**Fig. 2A**). Running is largely hard-wired in mice, but wild-type mice do become more adept at staying on the rod with training, so this seemed to present a suitable test for the improvement of a basic motor skill. Cumulatively, the HTT-SD (constitutively phosphorylated) mice fell off the rotarod more quickly than the wild-type mice (**Fig. 2B**); the HTT-SA mice showed a trend that did not quite reach significance. Looking at only the first trial on Day 1, there were no significant differences between genotypes (**Supplementary Fig. 2A**). Since mutant mice did not differ from WT in native motor coordination, strength, or resistance to prolonged activity in the open field, we examined motor skill learning on a daily basis over 80 sessions (10 sessions per day for 8 consecutive days). Over the eight days, the wild-type (WT) mice nearly doubled their latency to fall over the first three days/30 sessions of training (the learning phase) and then maintained that skill (the consolidation phase). The HTT-SD mice did not improve as much during the initial learning phase, however, and plateaued at a lower level of skill (**Fig. 2C**). The HTT-SA mice had an elongated learning curve but improved by the eighth day to wild-type level.

**Figure 2.**
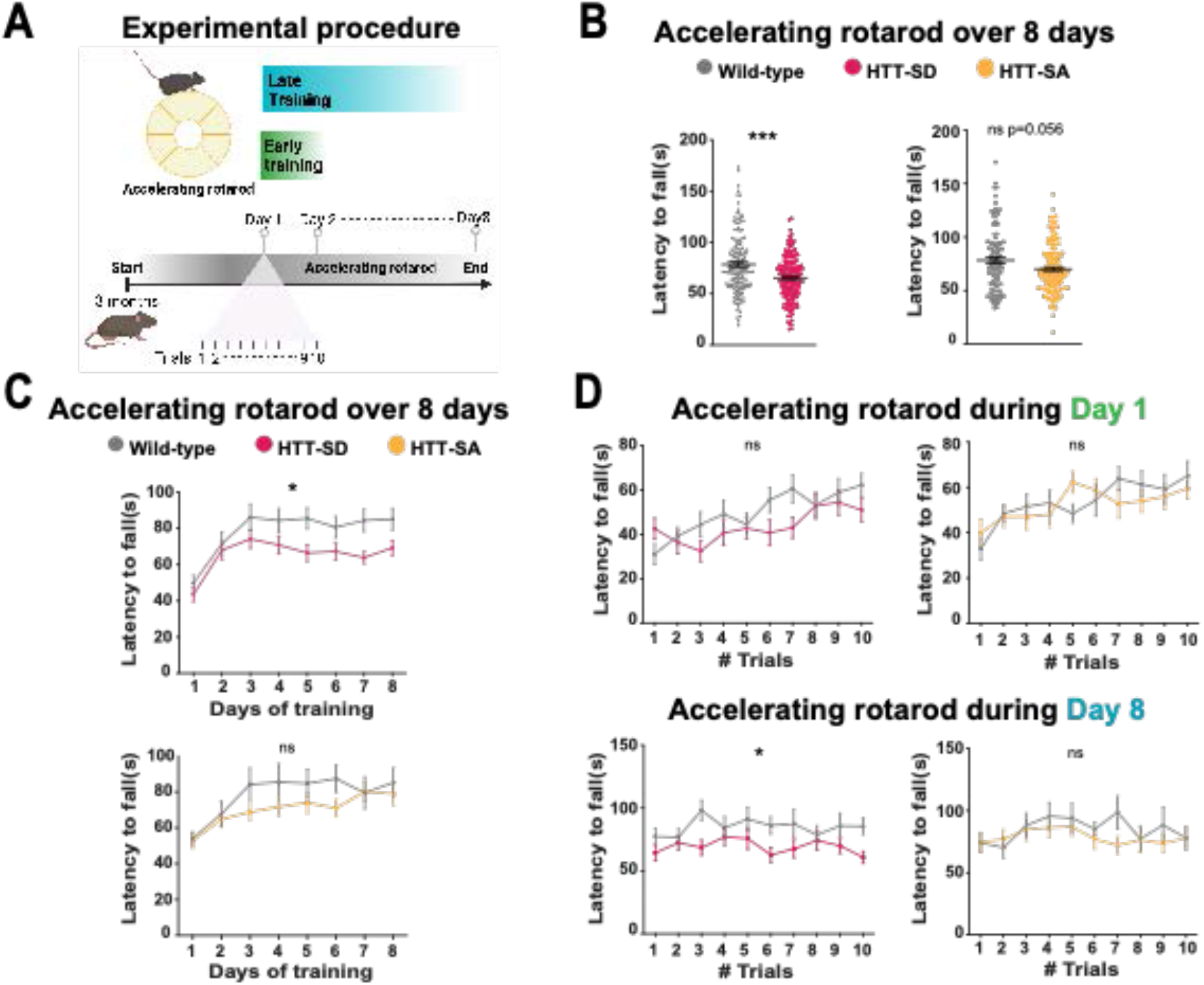
Constitutive phosphorylation of HTT at S421 impairs motor skill learning in mice. (**A**) Schematic of the accelerating rotarod protocol assessing the mouse motor skill learning over 8 days with 10 sessions per day. (**B**) Mean time to fall off the rotarod over 80 sessions (10 sessions per day for 8 days) for HTT-SD mice (*** p<0.001: Mann-Whitney test) and HTT-SA mice (ns=non significant: Mann-Whitney test). (**C-D**) Mean time to fall off the rotarod over 10 sessions (**C**) per day for HTT-SD mice (*p<0.01: Two way ANOVA) and HTT-SA mice (ns=non significant: Two way ANOVA) and (**D**) of the first and last day for HTT-SD (ns=non significant: Two-way ANOVA at day 1 and *p<0.01: Two-way ANOVA at day 8 followed by Sidak’s multiple comparisons test where p<0.01 at trial 3 and 10) and HTT-SA (ns=non significant: Two-way ANOVA at day 1 and 8). Histograms present the means +/-SEM of at least 3 cohorts and a total of N=20 WT, 20 HTT-SD and 13 WT, and 18 HTT-SA 3-month-old littermate male mice.

To better characterize this motor skill deficit, we focused on the first and the last days of learning. The first day of motor skill learning showed that HTT-SD had greater difficulty adjusting to the rod initially than WT mice but eventually improved (**Fig. 2D, Supplementary Fig. 2B**). On Day 8 the defect in motor skill learning was significant throughout most of the sessions (**Fig. 2D, Supplementary Fig. 2E**). HTT-SA mice showed a milder motor learning deficit with a less clear pattern. Because the HTT-SA neurons did not show significant changes in axonal transport or glutamate release (**Fig. 1B, C**), we focused thereafter on HTT-SD mice and neurons.

### HTT phosphorylation increases release probability and number of SVs at corticostriatal synapses

Motor skill learning relies on communication between the dorsal striatum and layer V pyramidal neurons in the motor cortex via the release of glutamate by the cortical afferences (Graybiel and Grafton, 2015; Jin and Costa, 2015; Perrin and Venance, 2019; Yin and Knowlton, 2006). We therefore performed whole-cell recordings of medium-sized spiny neurons (MSNs) located in the dorsolateral striatum (DLS) in acute corticostriatal brain slices from WT and HTT-SD adult mice (**Fig. 3A**; see Methods) and analyzed the spontaneous excitatory postsynaptic currents (sEPSCs). The HTT-SD mice did not differ from WT in sEPSC amplitude but did show greater sEPSC frequency (**Fig. 3B**).

**Figure 3.**
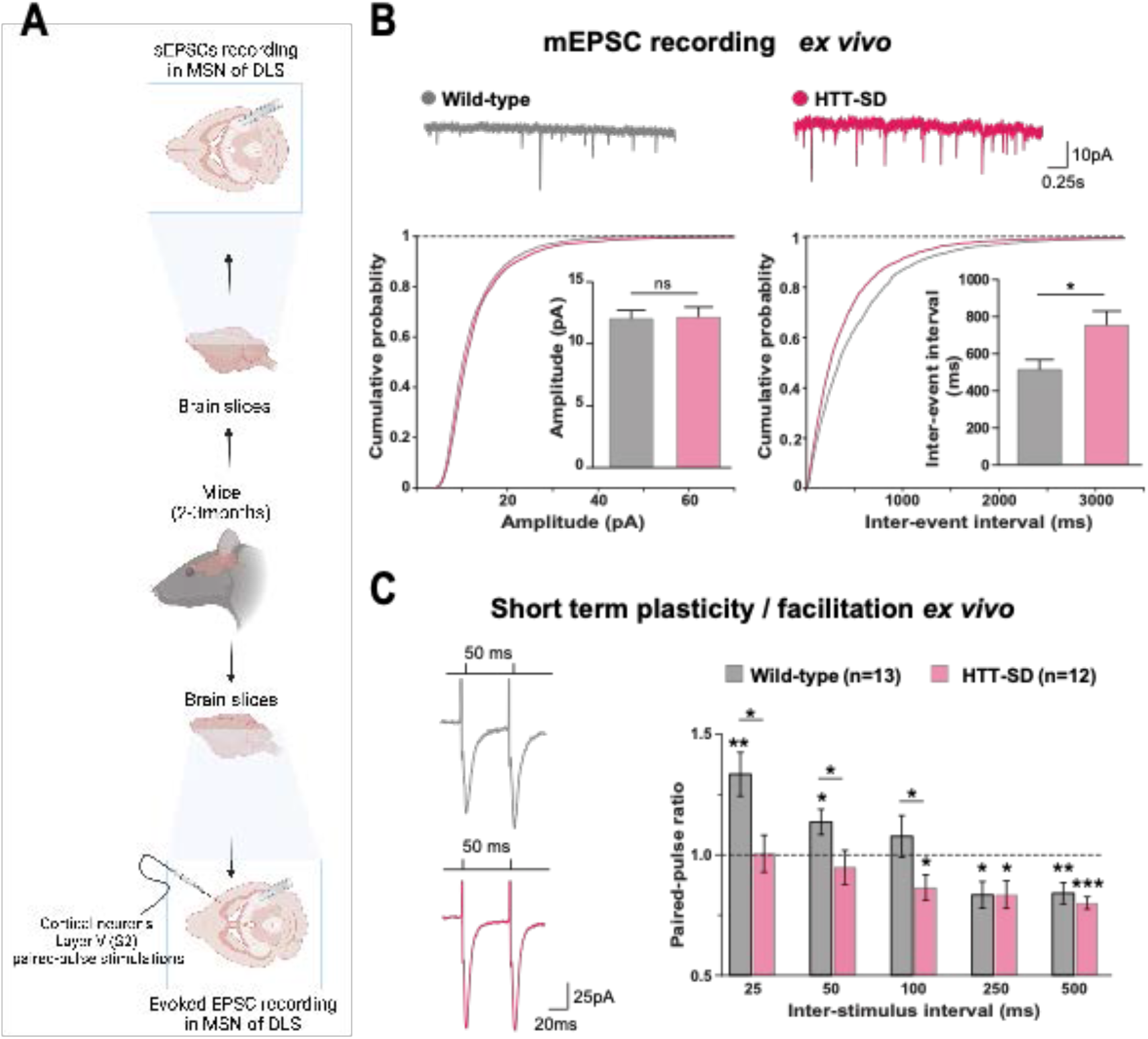
HTT phosphorylation at S421 increases the probability of release in the corticostriatal network *ex vivo*. (**A**) Schematic of the procedure for sEPSC recording (*top*) and paired-pulse stimulation (*bottom*) in MSNs within the DLS. (**B**) Representative traces, cumulative probability of the mean amplitude, and mean interevent intervals (p<0.05) of sPECS in MSNs within the DLS in 2-3-month-old WT and HTT-SD mice. Both parameters were recorded from WT (10 MSN) and HTT-SD mice (10 MSN). (**C**) Representative traces and quantification of the paired-pulse ratio per interstimulus interval of electrophysiological response of MSNs in the DLS after stimulation in S2 in 2-3-month-old WT and HTT-SD mice. In contrast to WT neurons, HTT-SD MSN response from 25 to 50 ms showed no facilitation (paired-pulse ratio=1) but depression at 100 ms (*p<0.05 for WT/HTT-SD at 25 ms, 50 ms, and 100 ms, WT at 50 ms, HTT-SD at 100 ms and WT and HTT-SD at 250 ms. **p<0.01 for WT at 25 ms and 500 ms. ***p<0.001 for HTT-SD at 500 ms). Paired pulse ratios were recorded from 13 WT MSNs and 12 HTT-SD MSNs from at least N=3 mice.

We next recorded EPSCs evoked by paired-pulse stimulations of layer V cortical neurons from the somatosensory S2 cortex and the corresponding corticostriatal projection field in the dorsal striatum at various interstimulus intervals (ISI: 25, 50, 100, 250, and 500 ms) to assess the probability of release at MSN corticostriatal synapses in WT and HTT-SD mice (**Fig. 3C**). Paired-pulse ratio (PPR) analysis revealed that in WT mice, corticostriatal short-term plasticity was facilitated for short ISIs (25 and 50 ms), as previously described (Goubard et al., 2011), but there was no detectable facilitation in HTT-SD mice (**Fig. 3C**). For 100 ms ISI, short-term depression was observed in HTT-SD but not in WT mice. For longer ISIs (250 and 500 ms), both genotypes showed similar depression.

The greater frequency of sEPSCs and lower facilitation in HTT-SD MSNs indicate that constitutively phosphorylated HTT augments the glutamate release probability of pyramidal cells. These findings are in agreement with the greater number of exocytic events in HTT-SD neurons within microfluidic devices. Since the increased frequency of sEPSCs and decreased facilitation indicate changes in the number of synaptic vesicles at the presynapse (Park et al., 2012; Pulido and Marty, 2017), we investigated the number of SVs at axon terminals within the corticostriatal network using electron microscopy. We focused on synapses formed between cortical neurons from the somatosensory cortex connecting with neurons from the DLS. According to the morphology of both the spines and the synapses, we counted the number of SVs in glutamatergic afferences within the DLS. We observed a greater number of SVs at the axon terminals in the brain of HTT-SD mice than in WT mice for the same number of synapses (**Fig. 4A**). We conclude that constitutive HTT phosphorylation favors the anterograde transport of SVPs, leading to a greater number of SVs at synapses in vivo.

**Figure 4.**
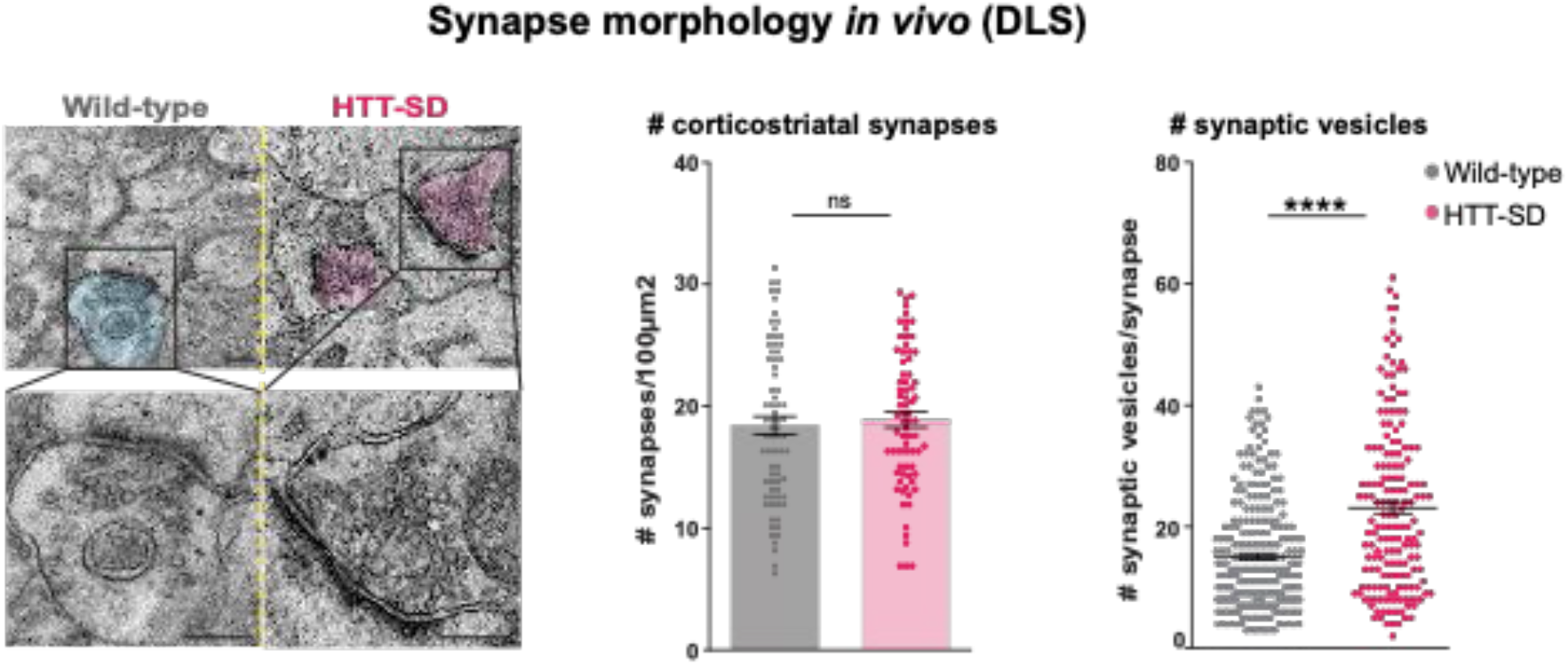
HTT phosphorylation increases the number of SVs at corticostriatal synapses. Representative images of SVs at the corticostriatal synapse were obtained by electronic microscopy from DLS of 3-month-old male WT and HTT-SD mice (left). Scale = 200 nm. *Middle:* The number of synapses at the corticostriatal synapse per 100 µm^2^ in DLS in 3 WT and 3 HTT-SD brains of littermate mice on N=74 WT striatal areas and 74 HTT-SD striatal areas. *Right:* The number of SVs at the corticostriatal synapse in N=5 WT and 3 HTT-SD mouse brains on N=279 WT and 171 HTT-SD axon terminals (**** p < 0.0001) (right). Histograms are represented with the means +/-SEM. Significance was determined using the Mann-Whitney test.

### HTT recruits KIF1A to vesicles

The anterograde transport of SVP is driven predominantly by the kinesin-3 motor KIF1A (Guedes-Dias and Holzbaur, 2019). HTT and KIF1A interactomes suggested a possible interaction between the two proteins (Shirasaki et al., 2012; Stucchi et al., 2018), but HTT has not yet been shown to interact functionally with KIF1A. We observed KIF1A in the proteome of HTT-associated vesicles (**Fig. 5A**)(Migazzi et al., 2021). We found that KIF1A colocalizes with HTT immunopositive puncta in free-cultured cortical neurons at DIV5 using a two-dimensional stimulated emission depletion (2D-STED) super-resolution microscope (**Fig. 5B**). We confirmed this observation after permeabilization of isolated axons within distal part of microfluidics axonal compartments and found an increased colocalization of KIF1A with HTT in HTT-SD neuronal network as compared to WT circuit (**Fig. 5C**). We next prepared vesicular-enriched fractions from WT and HTT-SD mouse brains and immunoblotted them for KIF1A. The vesicular/cytosolic ratio was greater for KIF1A in HTT-SD (**Fig. 5D** and **Supplementary Fig. 3A**), while p150 remained constant and the cellular levels of KIF1A did not differ between WT and HTT-SD mice (**Fig. 3E** and **Supplementary Fig. 3A**). KIF1A and HTT thus colocalize on VAMP2-positive vesicles, and S421 phosphorylation causes more KIF1A to be recruited to the vesicles.

**Figure 5.**
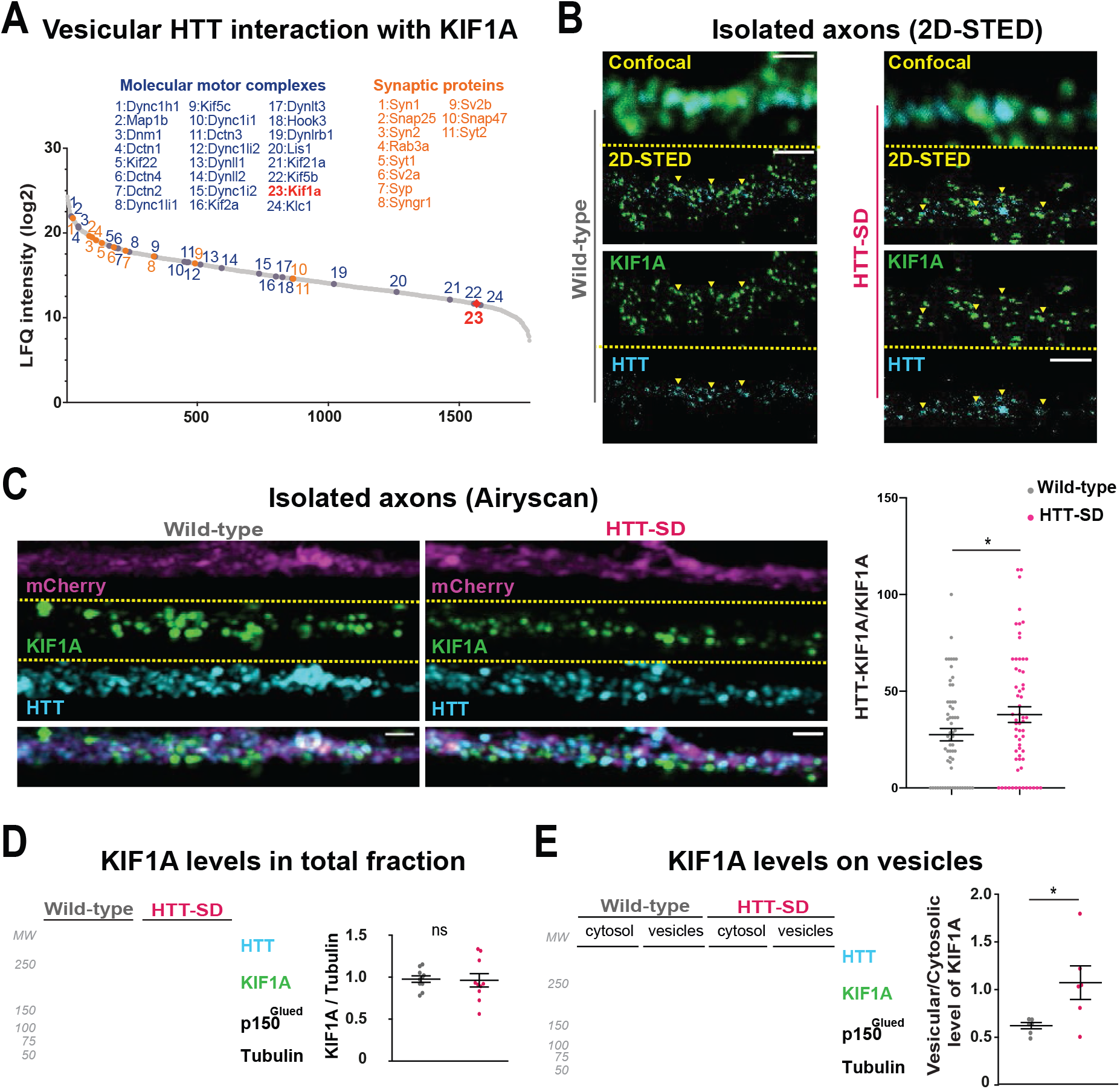
HTT phosphorylation recruits KIF1A on VAMP2-mCherry vesicles. (**A**) Mass spectrometry analysis of vesicles purified from mouse brains shows KIF1A among HTT-associated vesicular proteins. Gray: identified proteins, blue: motor complex, orange: synaptic proteins. (**B**). Confocal and 2D-STED images of free-cultured neurons at DIV5 showing the colocalization of KIF1A and HTT. Scale bar: 1*μ*m. (**C**) Representative immunofluorescence labeling revealing HTT (cyan), KIF1A (green), and VAMP2-mCherry (magenta) within cortical axons localized in long channels of the microfluidics. The images were acquired in a specific region of interest and processed by an Airyscan detector (Scale bar: 1 µm). The distribution analysis shows that there is greater colocalization of HTT and KIF1A on KIF1A^+^ vesicles in the HTT-SD condition than in WT. The graph represents means +/-SEM of 3 independent experiments reproducing a corticostriatal network of WT or HTT-SD neurons in at least 3 microfluidics devices per experiment. Significance was determined using the Mann-Whitney test (* p<0.05; N=61). (**D-E**) Western blot analysis for HTT, KIF1A (both bands), p150^Glued,^ and tubulin from total (**D**) and vesicular (**E**) fractions of N=6 WT and 6 HTT-SD brains. Significance was determined using the Mann-Whitney test (* p < 0.05). **Figure 5—source data 1** Western blot scans for the data presented in Figure 5d (KIF1A levels in whole brain lysates). Shown in red are the cropped regions presented in Figure 5d. Films containing the first batch of samples (Gel 1) are shown. **Figure 5—source data 2** Western blot scans for the data presented in Figure 5e (KIF1A levels in brain vesicular fractions). Shown in red are the cropped regions presented in Figure 5e. Films containing the second batch of samples (Gel 2) are shown.

### HTT-SD-mediated SVP transport depends on KIF1A

We next asked whether the phospho-HTT-mediated increase in SVP anterograde transport depends on KIF1A by using a validated sh-KIF1A (Kevenaar et al., 2016). Lentiviral expression of sh-KIF1A in cortical neurons led to an ∼83% decrease in the expression of KIF1A (**Supplementary Fig. 3B**). We then treated corticostriatal projecting neurons plated in microfluidic devices with lentiviruses expressing either a sh-scramble-GFP (sh-Scr) or the sh-KIF1A-GFP. We recorded axonal transport of VAMP2-mCherry vesicles at DIV12 and generated kymographs as before (**Fig. 6A**). We found that silencing KIF1A in WT cortical neurons decreased VAMP2 anterograde vesicle velocity, the number of anterograde vesicles (**Fig. 6B** and **Supplementary Fig. 3C**), the linear flow (**Supplementary Fig. 3C**) and the net directional flux of VAMP2 vesicles toward the axon terminals (**Fig 3E**).

**Figure 6.**
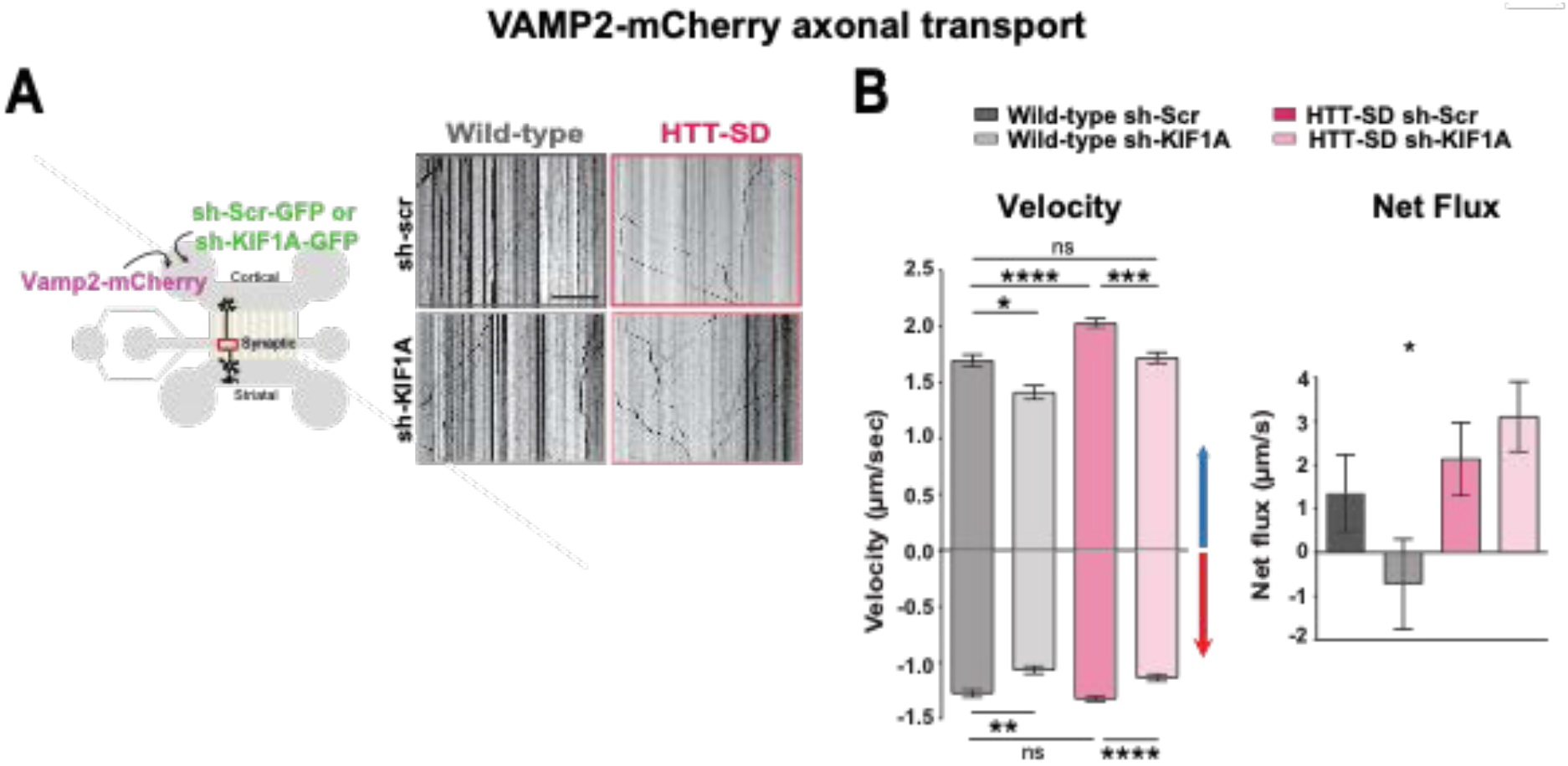
HTT-dependent axonal transport of SVPs is mediated by KIF1A. (**A**) Schematic of the 3-compartment microfluidic device with an indication of lentiviral transduction of VAMP2-mCherry and sh-scramble (sh-Scr-GFP) or sh-KIF1A (sh-KIF1A-GFP) lentiviruses at DIV8. On the right, representative kymographs of VAMP2-mCherry vesicle transport in axons for each condition. Scale bar = 25 µm. (**B**) Segmental anterograde and retrograde velocities (anterograde: * p<0.05, ***p<0.001, **** p < 0.0001; WT sh-Scr: 548 vesicles, N=WT sh-KIFA: 318 vesicles, DDsh-Scr:1129 vesicles, DD sh-Scr: 628 vesicles) (Retrograde: *p<0.05, **p<0.01, ****p<0.0001; N=WT sh-Scr: 583 vesicles, WT sh-KIFA: 396 vesicles, DD sh-Scr:1282 vesicles, DD sh-Scr: 620 vesicles) and directional net flux (* p<0.01; WT sh-Scr: 79 axons, N=WT sh-KIFA: 59 axons, DD sh-Scr:112 axons, DD sh-Scr: 89 axons; one-way ANOVA test) of VAMP2-mCherry vesicles in WT and HTT-SD neurons transduced with sh-Scr or sh-KIF1A lentiviruses. Histograms represent means +/-SEM of 3 independent experiments. Significance was determined using a one-way ANOVA followed by a Dunn’s multiple comparison test.

We next measured VAMP2 transport in HTT-SD neurons and found greater anterograde velocity, number of anterograde vesicles, and positive net directional flux (and/or linear flow) than in WT neurons (**Fig. 6B** and **Supplementary Fig. 3C**) thus confirming our previous results (**Fig. 1B**). Silencing KIF1A in HTT-SD reduced the anterograde velocity of VAMP2 vesicles close to values we observed in WT neurons (**Fig. 6B**). KIF1A silencing also reduced the number of anterograde vesicles (**Supplementary Figure 3C**), linear flow, and net directional flux in HTT-SD to values found in WT (**Fig 6B** and **Supplementary Fig. 3C**). The velocity and number of retrograde-moving VAMP2 vesicles was also lower in HTT-SD neurons (**Fig 6B**). This attenuation of retrograde transport might be linked to KIF1A’s reported role as a dynein activator (Chen et al., 2019).

We considered the possibility that the observed increase in SV release might be due in part to a synergistic action of BDNF at synapses, both because of the prominent role of HTT and its phosphorylation in regulating BDNF transport (Colin et al., 2008; Ehinger et al., 2020; Gauthier et al., 2004) and because synaptic BDNF levels regulate synaptic plasticity and SV release (Gangarossa et al., 2020; Park and Poo, 2013; Park et al., 2014; Tyler et al., 2006; Walz et al., 2006). Furthermore, dense-core vesicles (DCVs), including those containing BDNF, can be transported by kinesin-3 (Hung and Coleman, 2016; Lim et al., 2017; Lo et al., 2011; Stucchi et al., 2018). We therefore silenced KIF1A in cortical axons and measured BDNF-mCherry axonal transport in the distal part of axons at DIV12 (**Supplementary Fig. 4A**). Chronic HTT phosphorylation increased the anterograde transport of BDNF-containing vesicles, the linear flow as well as the net directional flux of BDNF vesicles in agreement with previous studies (Colin et al., 2008; Ehinger et al., 2020) (**Supplementary Fig. 4B**). Silencing KIF1A did not affect BDNF dynamics either in WT or in HTT-SD neurons. This indicates that the phospho-HTT-dependent increase in SVP anterograde transport, in contrast to BDNF, is mediated by KIF1A.

### HTT-KIF1A-mediated transport regulates the number of SVs at synapses

We next investigated whether SVP anterograde transport via the HTT-KIF1A complex regulates the number of vesicles at synapses. We injected lentiviruses encoding either sh-scramble-GFP or sh-KIF1A-GFP in layer V of the HTT-SD motor cortex (**Fig. 7A**), whose neurons project mainly to the DLS (Hunnicutt et al., 2016). We then counted the number of SVs at corticostriatal synapses from sections prepared from WT and HTT-SD brains injected with lentiviral sh-Scr or sh-KIF1A. WT sh-KIF1A presynapses showed significantly fewer SVs than WT sh-Scr presynapses (**Fig. 7B**). As previously shown (**Fig. 1B**), there were significantly more SVs at presynapses in HTT-SD, but this number reverted to WT levels in HTT-SD brains treated with sh-KIF1A (**Fig. 7C**). These data demonstrate that decreasing the phospho-HTT-mediated anterograde transport of SVPs by reducing KIF1A levels in corticostriatal projecting neurons reduces the number of synaptic vesicles at presynapses and suggests a tight link between increased axonal transport and synaptic SV content.

**Figure 7.**
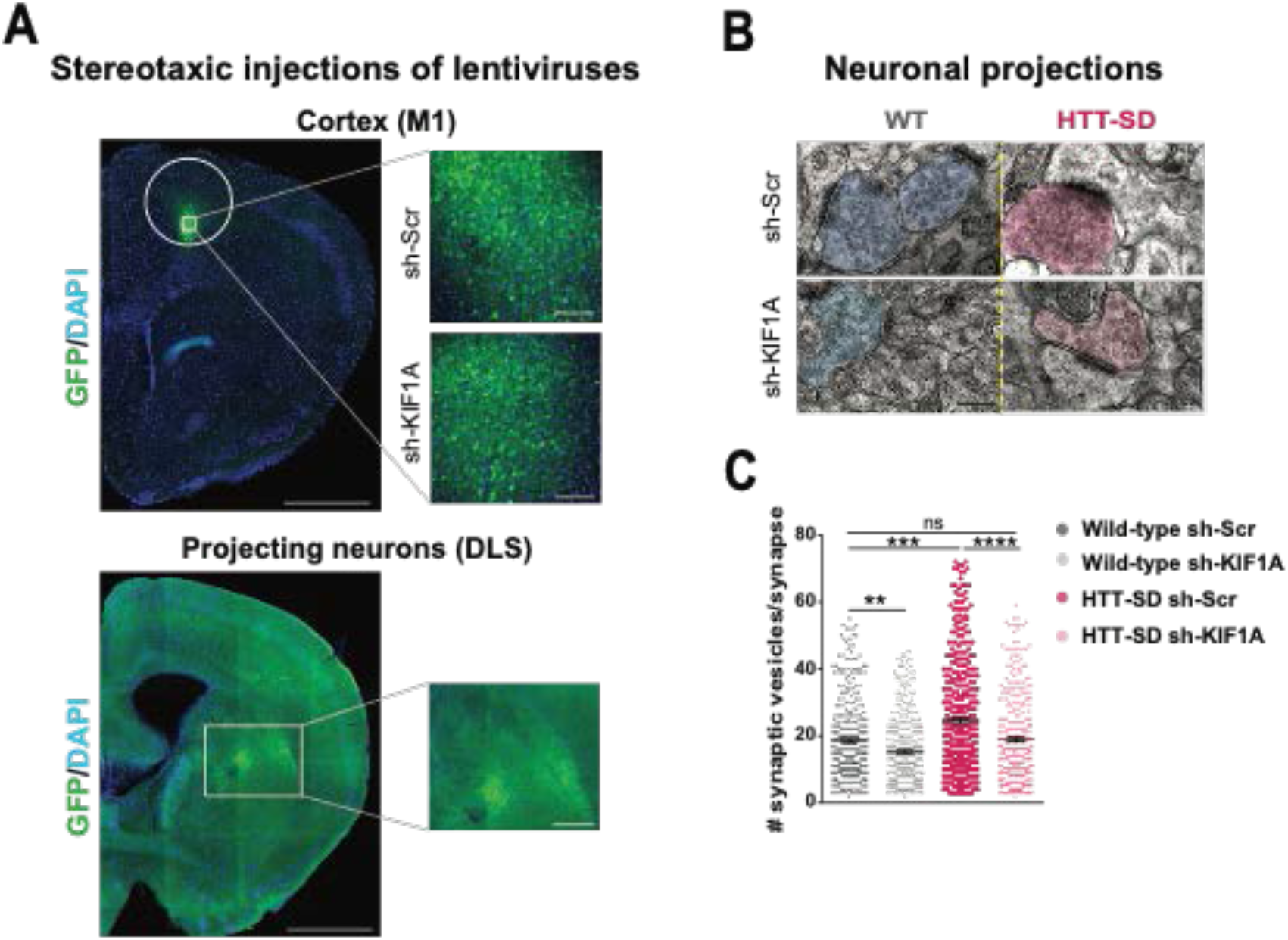
*In vivo* KIF1A silencing in mice restores the SV synaptic pool. (**A**) Immunolabeling of GFP within the injection site on a slice located at 1.5 mm before the bregma (left). Scale = 1 cm (insets, 100 µm). Immunolabeling of GFP within the projection site on a slice located at -0.3 mm after the bregma (right). Scale = 1 cm (inset, 250 µm). Nuclei are labeled with DAPI. (**B**) Representative images from electron microscopy of corticostriatal synapses and (**C**) quantification of the number of SVs at the corticostriatal synapse of 3 WT male mice injected with either sh-Scr or sh-KIF1A and 3 HTT-SD mice injected with sh-Scr or sh-KIF1A (** p < 0.01, *** p < 0.001, **** p < 0.0001; WT sh-Scr: 360 synapses, N=WT sh-KIF1A: 324 synapses, HTT-SD sh-Scr: 417 synapses, HTT-SD sh-KIF1A: 337 synapses). Scale = 200 nm. Histograms represent means +/-SEM. Significance was determined using a one-way ANOVA followed by a Dunn’s post-hoc analysis.

### HTT-KIF1A-mediated axonal transport of SVPs in corticostriatal projecting neurons regulates motor skill learning

To determine whether the modification in anterograde transport via the HTT-KIF1A complex within corticostriatal projecting neurons is responsible for the defect in motor skill learning we observed in HTT-SD mice, we injected lentiviral vectors encoding sh-Scr-GFP and sh-KIF1A-GFP into 3-month-old WT and HTT-SD mice. Three weeks after lentiviral injection we performed the same behavioral protocol as in Figure 1 (**Fig. 8A**). As with the non-injected mice (**Figure 2B**), HTT-SD mice did not show improvement over 8 days and had a much shorter latency to fall than WT mice (**Fig. 8B**). Silencing KIF1A improved the performance of the HTT-SD mice, especially over the first 5 days (**Fig. 8B**, left graph). A careful analysis of behavioral performance on the first and last days revealed that the improvement in motor learning of the HTT-SD mice via sh-KIF1A silencing was significant on the first day of training, but the effect did not last until the eighth day (**Fig. 8C**). Nonetheless, these findings indicate that HTT-KIF1A-mediated axonal transport of SVPs in the corticostriatal projecting neurons, a process modulated by phosphorylation, influences the number of SVs at synapses, the probability of release, and the efficacy motor skill learning.

**Figure 8.**
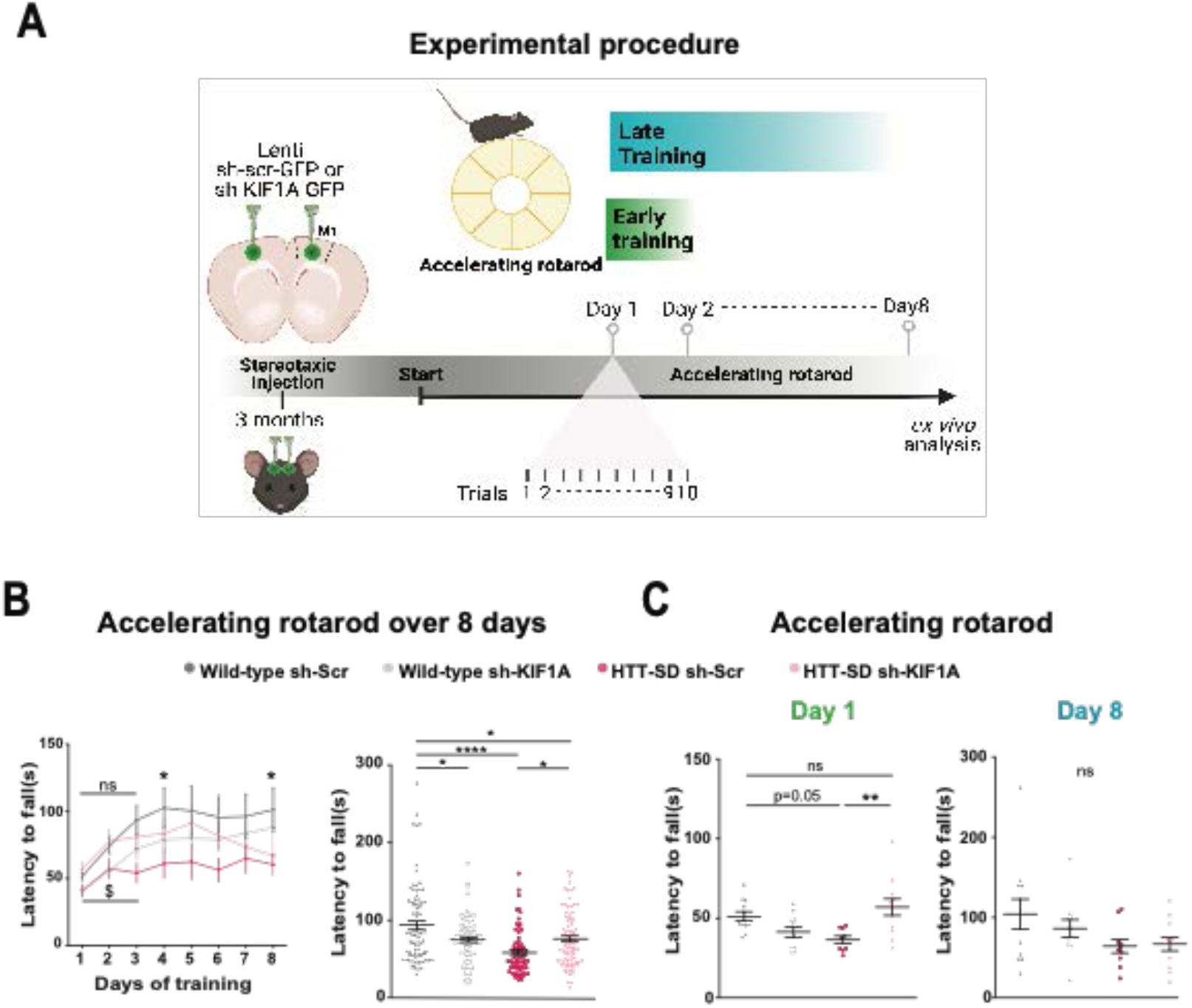
Motor skill learning defects of S421D mice are rescued by KIF1A silencing *in vivo*. (**A**) Schematic representation of the experimental procedure consisting in bilateral stereotaxic injections sites in the mouse brain followed by the accelerating rotarod protocol over 8 days. (**B**) *Left*: Mean time to fall off the rotarod over 8 days with 10 sessions per day. Two-way ANOVA comparing the four conditions showed significant differences between genotypes and silencing conditions (****p<0.0001). Tukey’s post-hoc analysis revealed significant differences between N=WT sh-Scr and HTT-SD sh-Scr at day 4 and day 8 (* p<0.05). In particular, although HTT-SD sh-Scr mice learned differently from WT sh-Scr from day 1 to 3 (^$^p<0.05: Two-way ANOVA), there were no differences between WT sh-scr and HTT-SD sh-KIF1A conditions, ns = not significant. *Right*: Mean time to fall off the rotarod during the 8 days (*p<0.05; ****p<0.0001). (**C**) Mean time to fall off the rotarod the first day (**p<0.01) (*left*) and the last day (ns=non significant, *right*), for the four conditions. Histograms are represented with the means +/-SEM of at least 3 cohorts containing N=12 WT sh-Scr, 11 WT sh-KIF1A, 10 HTT-SD sh-Scr, and 12 HTT-SD sh-KIF1A 3-month-old littermate male mice. Significance was determined using a two-way ANOVA test followed by Tukey’s post hoc analysis.

## Discussion

We show here that HTT acts as a scaffolding protein for KIF1A, thereby regulating the transport of SVPs and by extension influencing synaptic function. Specifically, HTT’s phosphorylation status fine-tunes SVP transport efficiency. Genetically blocking phosphorylation or dephosphorylation at S421 impaired motor learning, and abolishing KIF1A activity in the context of constitutive phosphorylation only partially restored motor performance on the rotarod. Such a genetic approach is rather blunt compared to the sensitivity of (de)phosphorylation in responding to cellular signals, yet it enabled us to answer the question that motivated the study and show that axonal transport does influence synaptic homeostasis, with consequences for circuit function and behavior.

### Huntingtin and the regulation of SVP axonal transport

This work places HTT among the proteins that participate in SVP transport (Guedes-Dias and Holzbaur, 2019) and closes a loop opened by the discovery that DENN/MADD, a Rab3-GEP that binds to KIF1A (and KIF1Bß) regulates SVP binding to microtubules according to Rab3’s nucleotide state (Niwa et al., 2008). Rab3 is part of the HTT interactome (Shirasaki et al., 2012) and enriched in SVs (Takamori et al., 2006); previous work in Drosophila larval axons showed that reducing HTT levels decreases the transport of Rab3-positive vesicles (White et al., 2015). In the context of Huntington’s disease, both Rab3 levels and the conversion from GTP to GDP state are dysregulated, which is consistent with a role for HTT in the transport of SVPs.

Studies in Huntington’s disease models have revealed alterations in HTT phosphorylation at S421 that could result from defects in Akt, the S421 kinase, or dysregulation of the phosphatases Calcineurin (PP2B) and PP2A (Humbert et al., 2002; Metzler et al., 2010; Pardo et al., 2006; Warby et al., 2005). Whether SVP trafficking is also altered and whether the specific restoration of SVP transport through HTT phosphorylation could ameliorate pathology need to be investigated. That such a study would be worthwhile is suggested by the fact that promoting HTT phosphorylation is neuroprotective, as it restores the transport and release of BDNF transport and release (Humbert et al., 2002; Kratter et al., 2016; Pardo et al., 2006; Pineda et al., 2009; Warby et al., 2009; Zala et al., 2008). It is interesting to note in this context that stimulation of glutamatergic corticostriatal connections in HD reverses motor symptoms in HD mice (Fernandez-Garcia et al., 2020), suggesting that reestablishment of synaptic release capacities could have therapeutic potential.

### The fine-tuning of SVP transport regulates synapse homeostasis and proper neurotransmission

Several studies have linked a reduction in axonal anterograde transport of SVPs to synapse function. Indeed, genetically impairing KIF1A reduces the number of SVs at nerve terminals and causes postnatal lethality (Yonekawa et al., 1998). KIF1A loss-of-function variants, most of them located within the conserved motor domain, reduce SVP transport and are associated with four diseases: autosomal recessive hereditary sensory neuropathy IIC, autosomal dominant mental retardation (ADMR) type 9, autosomal recessive spastic paraplegia (SPG) type 30, and autosomal dominant herditary spastic paraplegia (HSP) (Pennings et al., 2020). That all these disorders involve lower extremity spasticity and weakness reflects the extra challenge presented by the extremely long axons of the peripheral nervous system, but the cognitive deficits in ADMR type 9 show that disruptions in axonal transport can clearly disturb synaptic transmission and synaptic strength, with obvious consequences for learning and memory (Guedes-Dias et al., 2019; Zhang et al., 2017).

We found that increasing axonal transport of SVP via phospho-HTT-mediated KIF1A activation is equally problematic: it increases the number of SVs in the synaptic pool to the detiment of synaptic function and motor skill learning. This is consistent with the fact that the KIF1A gain-of-function mutation V8M, which causes another type of HSP, leads to greater anterograde transport of vesicles and abnormal accumulation of vesicles at synapses (Chiba et al., 2019; Gabrych et al., 2019). Further evidence for the sensitivity of synapses to both too little and too much SVPs comes from point mutations in the autoinhibitory domain of *unc-104* that cause hyperactive axonal transport and abnormal accumulation of synaptic vesicles (Cong et al., 2021). Similarly, depletion of kinesin-binding protein, KBP, which inhibits KIF1A activity, leads to the abnormal accumulation of both KIF1A and vesicles at neurite terminals (Kevenaar et al., 2016), and nonsense mutations of KBP cause Goldberg-Shprintzen syndrome (GOSHS), which is characterized by intellectual disability, microcephaly, and axonal neuropathy (Dafsari et al., 2015; Valence et al., 2013).

Together, these studies indicate that SVP transport is normally fine-tuned to ensure the proper quantity of SVs at the synapse and effective synaptic function.

### Axonal SVP transport, the synaptic vesicle pool, and SV release probability

Within the presynapse, SVs are organized into different pools—namely, the readily releasable pool (RRP), the recycling pool, and the reserve pool—according to their composition, age, and distance from the active zone (Crawford and Kavalali, 2015; Kaeser and Regehr, 2017; Rizzoli, 2014; Truckenbrodt et al., 2018). The RRP contains the SVs ready to be released upon neural activity and is thought to be refilled by SVs from the reserve pool by a slow process (hundreds of milliseconds) (Pulido and Marty, 2017; Rizzoli, 2014). The phenomenon of short-term facilitation is thought to compensate for the SV depletion of the RRP by increasing SV docking after a first stimulation (Pulido and Marty, 2017). We hypothesize that in our HTT-SD mice, the larger number of SVs at the presynapse increased the number of docked SVs and SVs released during the first stimuli, which could impair the increase in SV docking normally observed during facilitation.

### The dorsolateral striatum and motor skill learning

We found that lowering KIF1A levels within M1 neurons that project mainly to the DLS rescued motor skill learning, a process that is usually attributed to the DMS area (Costa et al., 2004; Yin et al., 2009). Recent studies have suggested that both DMS and DLS could be engaged during learning (Bergstrom et al., 2018; Gremel and Costa, 2013; Kimchi et al., 2009; Kupferschmidt et al., 2017; Perrin and Venance, 2019; Stalnaker et al., 2010; Thorn et al., 2010), so targeting DLS might be sufficient to rescue the development of motor skill during the first days of rotarod training. This would not, however, explain the lack of rescue upon KIF1A silencing in HTT-SD mice during the consolidation phase, when WT mice show that they maintain the skill level that peaked at Day 3. Rather, this could be related to the observation that during consolidation, the number of neurons that are activated upon stimulation of the DLS falls as the circuit streamlines its connections (Badreddine et al., 2022; Cao et al., 2015; Picard et al., 2013). In other words, as the motor skill is mastered, fewer neurons are needed to code the activity. Thus, it is possible that this small number of DLS connections responsible for consolidation of a learned skill would not have been targeted by the sh-KIF1A virus since it is unlikely that its expression would cover the entire DLS.

Altogether, our results highlight the importance of axonal SVP transport for synaptic transmission and identify a role for the HTT-KIF1A pathway as a regulator of SVP transport, synapse function, and motor skill learning within corticostriatal projecting neurons.

## Material and Methods

### Contact for Reagent and Resource Sharing

Further information and requests for resources and reagents should be directed to and will be fulfilled by the Lead Contact, Frédéric Saudou.

### Experimental Model and Subject Details Mice

*Hdh*^*S421A/S421A*^ and *Hdh*^*S421D/S421D*^ mice (referred to as HTT-SA and HTT-SD mice, respectively) have been previously described (Thion et al., 2015). They were generated by the Mouse Clinical Institute (Strasbourg, France). Briefly, these C57BL/6J mice were knocked-in with a point mutation replacing the serine 421 by an alanine or an aspartic acid, respectively. All mice were maintained with access to food and water *ad libitum* and kept at a constant temperature (19-22 °C) and humidity (40–50%) on a 12:12 hour light/dark cycle. All experimental procedures were performed in an authorized establishment (Grenoble Institut Neurosciences, INSERM U1216, license #B3851610008) in strict accordance with the directive of the European Community (63/2010/EU). The project was approved by the French Ethical Committee (Authorization number: APAFIS#18126-2018103018299125 v2) for care and use of laboratory animals and performed under the supervision of authorized investigators. For behavior studies, only males were used at 3-4 months of age. Behavioral studies compared littermates, homozygous (*Hdh*^*S421DorA/S421DorA*^) or wild type (*Hdh*^*+/+*^) mice. Those mice were then used for electrophysiological and biochemical studies or processed to be imaged by electron microscopy. The number of animals was limited to the minimum number necessary per group in order to have a chance of at least 80% of detecting a significant difference (power 1-β) and a risk of error α of 5%. This number was determined using a statistical test for estimating the optimal sample size using the variances determined in a preliminary study so as to reduce the number of animals used as much as possible while keeping enough to avoid compromising the validity of the experiments that was carried out.

For biochemistry and neuronal culture (E15.5), the sex distinction of homozygous or WT mice was not made. Specific ages used for each experiment are indicated in the figure legends. C57BL/6J mice, purchased from Charles River Laboratory, were used for backcrosses to maintain the colony and to obtain WT E15.5 pups.

### Primary neuron culture and transduction

Primary cortical and striatal neurons were dissected from E15.5 wild type (C57Bl/6J) or HTT-SA or HTT-SD mouse embryos as previously described (Liot et al., 2013). They underwent a chemical dissociation with papain cysteine solution, DNase (1/100), and FBS (1/10) and were finally mechanically dissociated. They were re-suspended in a growing medium containing a Neurobasal medium, 2% B27, 1% Penicillin/streptomycin, and 2 mM glutamax (5 × 10^6^ cells in 120 µl). Cortical neurons were plated in the presynaptic chamber coated with poly-D-lysine (0.1 mg/ml) and striatal neurons were plated in the postsynaptic chamber coated with poly-D-lysin and laminin (10 µg/ml) with a final density of ∼7000 cells/mm^2^. A growing medium was added to the synaptic chamber to equilibrate the flux. Neurons were left in the incubator for 2 hours and then all compartments were gently filled with growing medium. Neurons were cultured at 37°C in a 5% CO2 incubator for 10-12 days.

Between DIV0 and DIV4, cortical neurons within the presynaptic compartment were transduced with lentiviruses expressing VAMP2-mCherry, VGLUT1-pHluorin, or BDNF-mCherry. Neurons were washed out the next day. At DIV 8, cortical neurons were transduced with sh-KIF1A or sh-Scr lentiviruses.

### Accelerating rotarod

Motor skill learning was assessed using an accelerating rotarod (LE8305, BIOSEB). The tests were performed during the beginning of the light phase on male littermates housed in the same cages. Cages were transported to the experimental room at least 30 minutes before the tests to allow habituation of the mice to the room kept at a constant temperature (19-22 °C) and humidity (40–50%). The day preceding the test, the mice were acclimated to the rod by one session (1 minute at 4 rpm). Then, the accelerating rotarod assay was performed over 8 consecutive days with 10 sessions per day per mouse, increasing the speed from 4 rpm to 40 rpm over 5 minutes. Each trial was separated by at least a 15-minute resting period. The latency and the speed to fall off from the rotarod were recorded.

### Stereotaxic injections

3-month-old HTT-SD and WT male mice were anesthetized by inhalation of isoflurane associated with a mix of oxygen and room air (3-5% of isoflurane for induction and 1-2% in the mask). The mouse head was then shaved and placed within the stereotaxic frame. The skin was incised, and the skull was bilaterally drilled. The capillary was inserted slowly. We injected bilaterally (position AP:1,54 ML: + or -1,6 DV:-0,8) 500 nl of the diluted lentivirus (1/3 dilution in saline solution of the KIF1A shRNA or the Scr shRNA) at 0.5 µl/min speed using a nanoinjector. The capillary was slowly removed one minute after the end of the injection to prevent the leak of the injected solution. The skull was then washed with saline solution, the skin was sutured and 1 ml of NaCl 0.9% was injected subcutaneously. After surgery, mice were put alone in a warmed cage and monitored daily throughout recovery.

### Plasmids

The VAMP2-mCherry construct was a kind gift from T. Ryan’s laboratory. *Vglut1* cDNA sequence was amplified from an adult mouse brain. Its sequence from the 104^th^ amino acid to the end was cloned after the pHluorin sequence in a Smal site in a superecliptic pHluorin containing vector used in (Fernandez-Alfonso and Ryan, 2008). KIF1A shRNA construct (JL-35, target sequence: GACCGGACCTTCTACCAGT) has already been published (Kevenaar et al., 2016). It has been inserted in a EGFP-pSuper vector between NdeI and PstI sites. The control Scramble shRNA is a mouse universal scramble obtained from the scrambled order of HIF-1α nucleotides (Target Sequence: GGGTGAACTCACGTCAGAA)(Yu et al., 2004). It has been inserted in a EGFP-pSuper vector between NdeI and PstI sites. BDNF-mCherry construct and lentivirus, previously published (Hinckelmann et al., 2016) have been used for axonal transport experiments.

For lentivirus production, all the plasmids were cloned into a pSin vector (Drouet et al., 2009) by Gateway system (Life Technology) at the GIN virus production facility as described before (Bruyere et al., 2020). VAMP2-mCherry, sh-KIF1A, and sh-Scr lentiviruses were produced by the ENS Lyon Vectorology Facility.

### Microfluidic fabrication

Microfluidic devices were generated as previously described (Lenoir et al., 2021; Virlogeux et al., 2018). Briefly, we modified the size of the microchannels (3 μm width, 3 μm height, and 450 μm length) of polydimethylsiloxane microfluidic device (Taylor et al., 2005). After amplification and production, microfluidic devices were sealed on Iwaki boxes using plasma cleaner. The upper chamber was then coated with poly-D-lysin (0.1 mg/ml) and the lower chamber was coated with poly-D-lysin (0.1 mg/ml) and laminin (10 μg/ml). After overnight incubation at 4°C, microfluidic devices were washed 2 times with the growing medium. Microchambers were then placed in the incubator until neurons were plated.

### Videomicroscopy

Videorecording of neurons plated in microfluidic devices was performed at DIV 12. Before recordings, DIV 12 neurons in the microchamber were carefully inspected and selected based on the absence of cell contamination. For double transductions (with sh-RNA lentiviruses), the transport of mCherry-tagged cargo was analyzed within GFP-positive axons. Images were acquired every 200 ms for 1 minute on an inverted microscope (Axio Observer, Zeiss) with X63 oil-immersion objective (1.46NA) coupled to a spinning-disk confocal system (CSU-W1-T3; Yokogawa) connected to an electron-multiplying CCD (charge-coupled device) camera (ProEM+1024, Princeton Instrument) at 37 °C and 5% CO_2_.

For the study of the exocytosis events, images were acquired every 200 ms for 1min on an inverted microscope (Axio Observer, Zeiss) with X63 oil-immersion objective (1.46NA) coupled to a spinning-disk confocal system (CSU-W1-T3; Yokogawa) with TIRF microscopy (Nikon/Roper, Eclipse Ti) equipped with a camera Prime 95B sCMOS (Telelyne Photometrics) at 37 °C and 5% CO_2_. The same three fields per microchambers were acquired before and after a 4AP-bicuculline (respectively 2.5mM and 50µM) stimulation of the presynaptic chamber, four times in total (1 before and three after stimulation).

### Immunostaining

Neurons from the reconstituted corticostriatal network were fixed with a PFA/Sucrose solution (4%/4% in PBS) for 20 minutes at room temperature (RT). After three washes of PBS, neurons were incubated first with a blocking solution (BSA 1%, NGS 2%, Triton X-100 0.1%) and then with primary antibodies for KIF1A (Abcam, #ab180153, 1:100, rabbit), HTT (Millipore, #MAB2166, 1:500, mouse), and mCherry (Fisher Scientific, #16D7, 1:200, rat) overnight at 4°C. The next day, neurons were washed three times with PBS followed by one-hour incubation at RT of appropriate secondary antibodies (1:1000) and finally washed again three times with PBS. Images were acquired with a X63 oil-immersion objective (1.4 NA) using an inverted confocal microscope (LSM 710, Zeiss) coupled to an Airyscan detector. For 2D-STED microscopy, we used the Abberior kit containing the secondary antibodies (STAR RED anti mouse or rabbit, STAR ORANGE anti mouse or rabbit) and coverslips were mounted with the Abberior mount solid. Images were taken with a 100X oil-immersion objective (1.46 NA) using the Abberior 2D-STEDYCON upright confocal microscope.

For brain slices, brains were incubated in PFA 4% overnight and washed with PBS three times the next day. Then, brains were cut into 100 µm-thick slices using a vibratome. The slices were incubated with a blocking solution (0.3% triton, 10%NGS in PBS) for 2 hours at RT and then with antibody against GFP (Institut Curie, A-P-R#06) overnight at 4°C. The day after, the primary antibody was removed by 3 washes of PBS before incubating the slices with the associated secondary antibody and finally with 3 washes of PBS. Finally, slices were incubated with DAPI (1/4000) for 15 minutes, washed three times with PBS, mounted on Superfrost slides by using Dako Faramount Aqueous Mounting Medium solution and coverslips. The slices were acquired with a x10 objective (0.45 NA) using a slide scanner (AxioScan Z1, Zeiss) and with a x10 objective (0.3 NA) using an inverted confocal microscope (LSM 710, Zeiss) coupled to an Airyscan detector to improve signal-to-noise ratio and to increase the resolution.

### Western blotting

Cortical neurons were plated in free culture, transduced at DIV1 with sh-KIF1A or Sh-scr, and lysed at DIV5 in NetN buffer (20 mM Tris-HCl pH8, 120 mM NaCl, 1mM EDTA, 0.5% NP40) complemented with protease inhibitor cocktail (Roche).

A vesicular fraction from brains was prepared as described in (Hinckelmann et al., 2016). Briefly, brains were homogenized in lysis buffer (10mM HEPES-KOH, 175 mM L-aspartic acid, 65 mM taurine, 85 mM betaine, 25 mM glycine, 6.5 mM MgCl2, 5mM EGTA, 0.5 mM D-glucose, 1.5 mM CaCl2, 20 mM DTT pH 7.2, protease inhibitor from Roche) on ice with a glass potter and then with a 25G needle. Lysates were then centrifuged (12000 RPM) and the supernatant, considered as the total fraction, is then centrifuged (3000 RPM for 10 minutes). The resulting supernatant was centrifuged (12 000 RCF for 40 minutes). The supernatant was then ultracentrifuged (100 000g) to obtain the vesicular fraction (the pellet) and the cytosolic fraction (the supernatant).

All types of lysed samples were dosed by a Bradford reagent to quantify the protein concentration and then analyzed by SDS-PAGE transferred to PVDF membranes. Then, membranes were incubated for 45 minutes in a 5% BSA TBST (10mM Tris pH 8, 150 mM NaCl, 0.5% Tween 20) solution and incubated with primary antibodies against KIF1A (Abcam # ab180153, 1:5000), p150 (BD laboratories, # 612708, 1:1000), Tubulin (Sigma #T9026, 1:1000) at 4°C, overnight. The next day, membranes were washed at least three times with TBST and incubated with secondary antibodies conjugated with Horseradish Peroxidase (HRP) against mouse or rabbit (1:1000) for two hours at RT. Membranes were finally revealed with ECL (Thermo Scientific) after three washes of TBST.

### Electron microscopy

We anesthetized 3-to 4-month-old animals with 1ml/kg of Doléthal^®^ and perfused them transcardially with cold PBS followed by 2% paraformaldehyde 2% glutaraldehyde and 0,1M cacodylate cold solution. We removed brains from the skull and fixed them in a 0.1M phosphate buffer pH7.2 with 2% of glutaraldehyde and 2% of paraformaldehyde for 48 hours at 4°C before obtaining 2 mm-thick or 100µm-thick slices from a mold and a vibratome, respectively. A 1mm square piece of tissue was removed from the dorsolateral striatum; samples were then fixed again with the same solution for 72 hours, washed with phosphate buffer, and then post-fixed in a 0.1M phosphate buffer pH 7.2 with 1% Osmium tetroxide for 1 hour at 4°C. After extensive washes with water, samples were then stained with a solution of 1% uranyl acetate pH 4 in water for 1 hour at 4°C. They were further dehydrated through a gradient of ethanol (30%-60%-90% and three at 100%) and infiltrated with a solution of 1/1 epon/alcohol 100% for 1 hour and followed by several baths of fresh epon (Fukka) for 3 hours. The resin was then poured into capsules containing the samples, heated at 60°C for 72 hours for polymerization, and finally cut into ultrathin sections with an ultramicrotome (Leica). Sample sections were then post-stained with fresh solutions of 5% uranyl acetate and 0.4% of lead citrate, observed with a transmission electron microscope at 80 kV (JEOL 1200EX) and images were acquired with a digital camera (Veleta, SIS, Olympus). Analysis was performed with ImageJ and quantification of the number of synapses was performed on axon-free neuropil regions (Zhang et al., 2015).

### Brain slice preparation and whole-cell patch-clamp recordings

All experiments were performed in accordance with the guidelines of the local animal welfare committee (Center for Interdisciplinary Research in Biology Ethics Committee) and the EU (directive 2010/63/EU). We prepared horizontal brain slices containing the somatosensory S2 cortex and the corresponding corticostriatal projection field in the dorsal striatum from mice (2-3-months old) using a vibrating blade microtome (VT1200S, Leica Micosystems, Nussloch, Germany). Brains were sliced in a 5% CO2/95% O_2_-bubbled, ice-cold cutting solution containing (in mM): 125 NaCl, 2.5 KCl, 25 glucose, 25 NaHCO_3_, 1.25 NaH_2_PO_4_, 2 CaCl_2_, 1 MgCl_2,_ and 1 pyruvic acid, and then transferred into the same solution at 34°C for 60 minutes and then moved to room temperature.

For whole-cell patch-clamp recordings, borosilicate glass pipettes of 4-6 MΩ resistance contained (in mM): 105 K-gluconate, 30 KCl, 10 HEPES, 10 phosphocreatine, 4 ATP-Mg, 0.3 GTP-Na, 0.3 EGTA (adjusted to pH 7.35 with KOH). The composition of the extracellular solution was (in mM): 125 NaCl, 2.5 KCl, 25 glucose, 25 NaHCO_3_, 1.25 NaH_2_PO_4_, 2 CaCl_2_, 1 MgCl_2,_ and 10μM pyruvic acid, bubbled with 95% O_2_ and 5% CO_2_. Signals were amplified using EPC10-2 amplifiers (HEKA Elektronik, Lambrecht, Germany). All recordings were performed at 34°C using a temperature control system (Bath-controller V, Luigs & Neumann, Ratingen, Germany) and slices were continuously superfused at 2-3 ml/min with the extracellular solution. Slices were visualized on an Olympus BX51WI microscope (Olympus, Rungis, France) using a 4x/0.13 objective for the placement of the stimulating electrode and a 40x/0.80 water-immersion objective for localizing cells for whole-cell recordings. The series resistance was not compensated. Recordings were sampled at 10 kHz, using the Patchmaster v2×32 program (HEKA Elektronik).

For paired-pulse protocols, electrical stimulations were performed with a bipolar electrode (Phymep, Paris, France) placed in the layer V of the somatosensory S2 cortex. Electrical stimulations were monophasic at constant current (ISO-Flex stimulator, AMPI, Jerusalem, Israel). Currents were adjusted to evoke 50-200pA EPSCs. Repetitive control stimuli were applied at 0.1Hz. For each ISI, 20 successive EPSCs were individually measured and then averaged. Variation of input and series resistances above 20% led to the rejection of the experiment. Off-line analysis was performed with Fitmaster (Heka Elektronik). Statistical analysis was performed with Prism 5.02 software (San Diego, CA, USA). All results are expressed as mean ± SEM. Statistical significance was assessed in non-parametric Mann Whitney, one-sample t-tests using the indicated significance threshold (p).

### Mass spectrometry

This analysis follows that of (Migazzi et al., 2021). Briefly, vesicular fraction from brains obtained as described earlier was first pre-cleared for an hour at 4°C with protein A Sepharose beads (Sigma Aldrich-P9424) and then immunoprecipitated for 3 hours at 4°C by agarose beads preincubated with rabbit anti-HTT D7F7 antibody (Cell Signaling, Cat#5656). To remove the non-specific binding, the beads were washed three times with the lysis buffer and bound proteins are finally eluted with Laemmli buffer. The HTT corresponding band on the western blot was cut and analyzed. MS was performed with a LTQ Orbitrap XL mass spectrometer (Thermo Scientific), equipped with a nanoESI source (Proxeon). The top eight peaks in the mass spectra (Orbitrap; resolution, 60,000) were selected for fragmentation (CID; normalized collision energy, 35%; activation time, 30 ms, q-value, 0.25). Proteins were identified using the MaxQuant software package version 1.2.2.5 (MPI for Biochemistry, Germany) and UniProt database version 04/2013.

### Quantification and Statistical Analyses Transport analysis

Vesicle velocity, directional flow, and vesicle number were measured on 100 µm of neurite using KymoTool Box ImageJ plugin, as previously described (Virlogeux et al., 2018). Anterograde or retrograde speeds describe, respectively, the mean speed of anterograde or retrograde segmental movement of a vesicle. Static vesicles are those without any movement during the recording. Linear flow and directionality were calculated as in (Virlogeux et al., 2018).

### Electrophysiology analysis

For each ISI, 20 successive EPSCs were individually measured and then averaged. Variation of input and series resistances above 20% led to the rejection of the experiment. Off-line analysis was performed with Fitmaster (Heka Elektronik). Statistical analysis was performed with Prism 5.02 software (San Diego, CA, USA). All results are expressed as mean ± SEM. Statistical significance was assessed in non-parametric Mann Whitney, one-sample t-tests using the indicated significance threshold (p).

### Immunostaining

Immunostained vesicles in distal axons were quantified as previously shown (Scaramuzzino et al., 2022). Briefly, we used a customized macro for imageJ where the images are enhanced using a DoG filter adapted to the vesicle size. Masks are created on each channel using manual thresholding that is kept constant for each individual channel and replicates. Finally, the number of particles is automatically counted for the single and dual channels and expressed as the percentage of co-localization.

### Exocytosis events

The same three fields per microchamber were acquired before and after a 4AP-bicuculline (respectively 2.5mM and 50µM) stimulation of the presynaptic chamber, four times in total (1 before and three after stimulation). Recording of the amplitude and the number of exocytosis events were automatized. The number of events was expressed as follows:

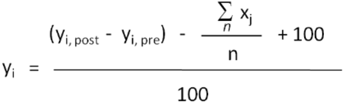

Where

- *y*_*i*_ expresses the difference of the number of events after (y_i, post_) minus before (y_i, pre_) stimulation in HTT-SD neurons of the field *i*,
- *x*_*j*_ expresses the difference of the number of events after minus before stimulation in WT neurons of the field *j* and *n* of them have been averaged.
- the final value was normalized to 1, *i*.*e*. a given HTT-SD neuron field, whose activity after stimulation increased as much as that of the average of the WT neuron fields, will display a value of 1.
- The amplitude of the signal from stimulated neurons was normalized by that of the same neuron before stimulation.

### Statistical analysis

Statistical calculations were performed using GraphPad Prism 6.0. Statistical parameters (Replication, sample size, SEM, etc…) are reported in the legend figure. For each data set, outliers were identified and removed from the analysis using the ROUT test (Q=1%). Then, the Shapiro-Wilk normality test with the threshold set at *α* = 0.05 was performed to assess the normality of the data.

According to whether the data set followed a normal repartition or not, parametric or non-parametric tests were performed respectively. Then, if two conditions were analyzed, a t-test (or a Mann-Whitney test if nonparametric) was used. If more than two conditions were compared, a one-way ANOVA followed by a Tukey ‘s post-hoc analysis (or a Kruskal-Wallis test followed by a Dunn’s post hoc analysis if nonparametric) was used. If the data set were dependent on another, a two-way ANOVA was used followed by Tukey ‘s post-hoc analysis if more than two groups are compared or a Sidak’s post-hoc analysis if only two groups are analyzed. For the non linear fit, the run test was performed to study whether the curve deviates systematically from the data. Low P value (ns) indicates that the curve poorly describes the data. *p < 0.05; **p < 0.01; ***p < 0.001; ****p < 0.0001; ns, not significant. N number corresponds to biological replicates.

## Data Availability

All data generated or analyzed during this study are included in the manuscript and supporting files. Source Data files have been provided for Figure 1, Figure 1-Figure Supplement 1, Figure 2, Figure 2-Figure Supplement 2, Figure 3, Figure 4, Figure 5, Figure 5-Figure Supplement 3, Figure 6, Figure 6-Figure Supplement 4, Figure 7, Figure 8.

## Acknowledgements

We thank Sebastien Carnicella, Sandrine Humbert, Alain Marty, members of the Saudou, Humbert, and Venance labs for comments; Vicky Brandt for critical editing; Béatrice Blot, Aurélie Genoux, Nagham Badreddine, Claire Seris and Camille Brodier for technical help; T.Ryan for the gift of pHluorin plasmid; C. Hoogenraad for the gift of sh-KIF1A plasmid; Y. Saoudi for help with image acquisitions and the Photonic Imaging Center of Grenoble Institut Neuroscience (PIC-GIN) which is part of the ISdV core facility and is certified by the *IBiSA* label. We acknowledge the contribution of Gisèle Froment, Didier Nègre and Caroline Costa and the AniRA lentivectors production facility from the CELPHEDIA Infrastructure and SFR Biosciences (UAR 3444/CNRS, US8/Inserm, ENS de Lyon, UCBL). This work was supported by grants from the European Research Council (ERC) under the European Union’s Horizon 2020 research and innovation program AdG grant agreement no. 834317, Fueling Tranport (F.S.); Agence Nationale de la Recherche: ANR-15-IDEX-02 NeuroCoG (F.S.) in the framework of the “Investissements d’avenir” program; ANR-18-CE16-0009-01 AXYON (F.S.); Fondation pour la Recherche Médicale, FRM, DEI20151234418 (F.S.) and AGEMED program from INSERM (F.S.). The Saudou laboratory is part of the Grenoble Center of Excellence in Neurodegeneration (GREEN). C.S. was supported by a Postdoctoral fellowship from: FRM (SPF20140129405) and EMBO LTF (ALTF 693-2015). H.V. was supported by a PhD fellowship from Association Huntington France and by a FRM fellowship (FDT201904008035).

## Author contributions

Conceptualization: HV, JB, CS, LV and FS; Methodology: HV, JB, HX, JB, YSA, and CS; Investigation: HV, JB, HX, YSA, and CS; Writing – Original Draft: HV, CS, and FS; Writing – Review & Editing: HV, JB, HX, JB, YSA, BD, CS, LV, and FS; Funding acquisition: LV, CS and FS; Supervision: JB, CS and FS

## Additional information

**Supplementary files** include 4 Supplementary Figures

---

**Figure S1.**
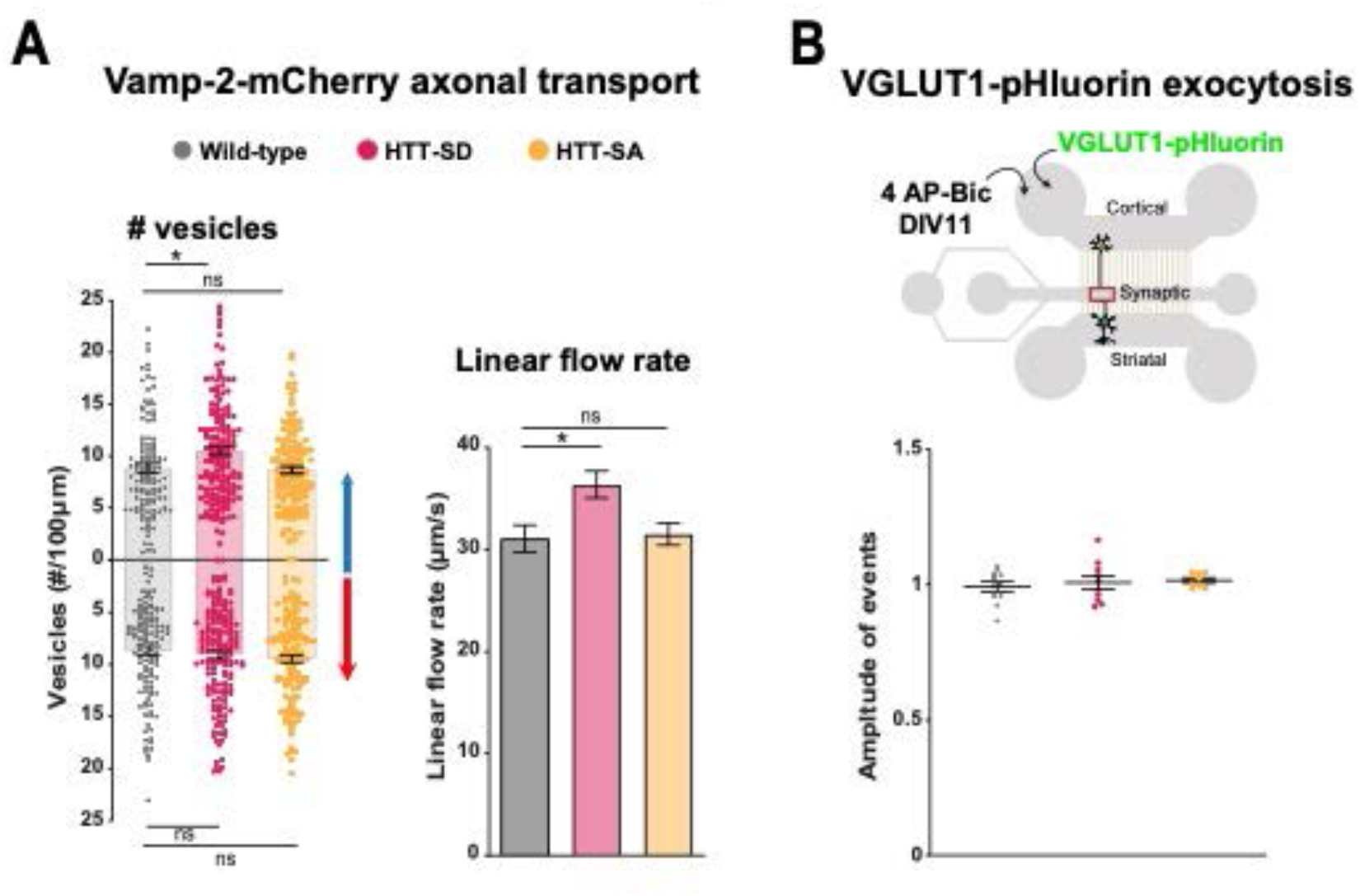
HTT phosphorylation at S421 increases SVP anterograde axonal transport and SV release. (**A**) Number of anterograde (*p<0.05; N=WT: 117 axons, HTT-SD: 156 axons and HTT-SA: 132 axons) and retrograde (*p<0.05; N=WT: 118 axons, HTT-SD: 159 axons and HTT-SA: 134 axons) VAMP2-mCherry axonal vesicles along 100 µm of axon in WT, HTT-SD and HTT-SA axons and their linear flow rate (*p<0.05; N=WT:118 axons, HTT-SD: 158 axons and HTT-SA: 133 axons). Histograms represent means +/-SEM of 3 independent experiments. Significance was determined using a one-way ANOVA followed by a Dunn’s multiple comparison test. (**B**) Representation of the 3-compartment microfluidic device with an indication of lentiviral transduction and stimulation with 4AP bicuculline. The amplitude of VGLUT-1 pHluorin exocytosis events within the synaptic chamber of the corticostriatal network was compared to that of non-stimulated condition (see Methods). Histograms represent means +/-SEM of 3 independent experiments. Significance was determined using one-way ANOVA followed by a Dunn’s multiple comparison test (ns=not significant).

**Figure S2.**
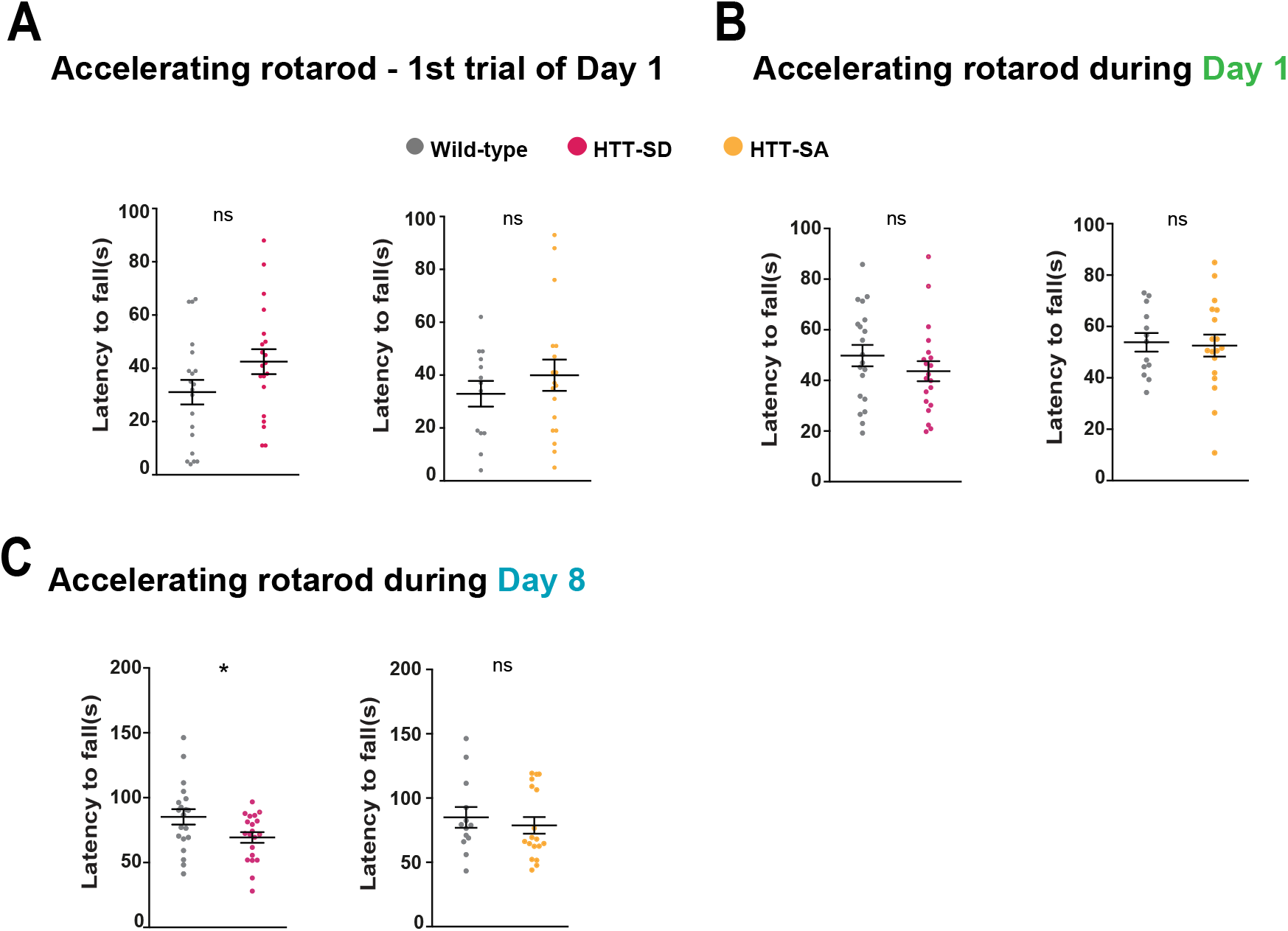
HTT phosphorylation at S421 impairs motor skill learning without affecting motor performance. (**A**) Mean time to fall off during the first trial of the first day for HTT-SD and HTT-SA mice. (**B**) Mean time to fall off during the first day and the last day (**C**) for HTT-SD (*p<0.05) and HTT-SA mice. Histograms present the means +/-SEM of at least 3 cohorts and a total of N=20 WT, 20 HTT-SD and 13 WT, and 18 HTT-SA 3-month-old littermate male mice.

**Figure S3.**
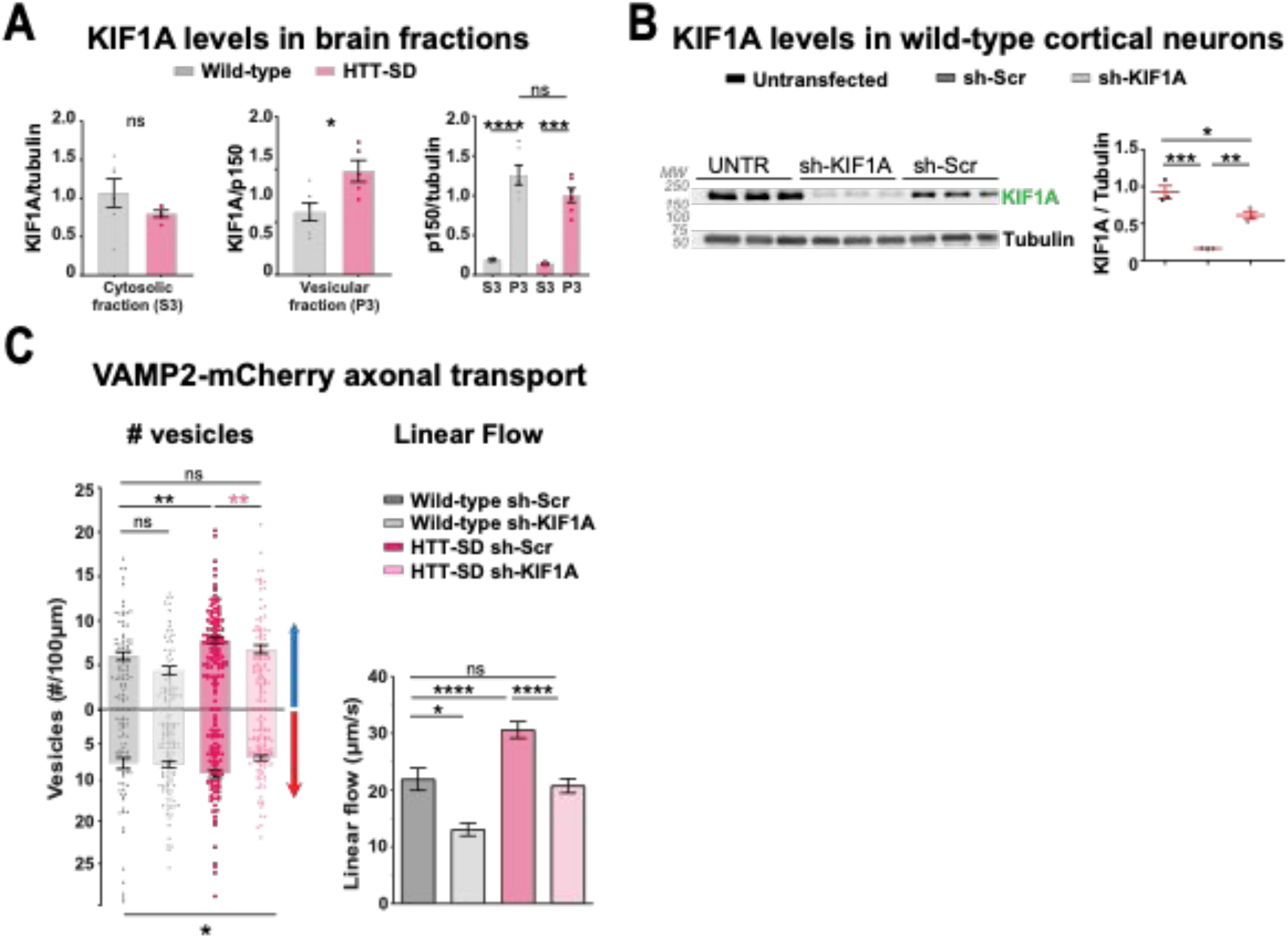
KIF1A levels in HTT-SD neurons regulate VAMP-2 axonal transport. (**A**) Western blot analysis of KIF1A levels (two bands) in the cytosol (left), the vesicles (p<0.05) (middle), and of p150^Glued^ in both fractions from N=6 WT and 6 HTT-SD brains. (**B**) Analysis of KIF1A levels by western blot in cortical neurons not-transduced and transduced with sh-KIF1A or sh-Scr lentiviruses (*p<0.05, **p<0.01, ***p<0.001; one way ANOVA followed by Tukey’s test). Histograms are represented with the means +/-SEM. (**C**) Number of anterograde (*p<0.05, **p<0.01; N=WT sh-Scr: 76 axons, WT sh-KIF1A: 59 axons, HTT-SD sh-Scr: 110 axons and HTT-SD sh-KIF1A: 86 axons) and retrograde (*p<0.05; N=WT sh-Scr: 60 axons, WT sh-KIF1A: 114 axons, HTT-SD sh-Scr: 79 axons and HTT-SD sh-KIF1A: 95 axons) VAMP2-mCherry axonal vesicles along 100 µm of axon in WT and HTT-SD and their linear flow rate (*p<0.05; N=WT sh-Scr: 75 axons, WT sh-KIF1A: 59 axons, HTT-SD sh-Scr: 107 axons and HTT-SD sh-KIF1A: 85 axons. One way ANOVA followed by Dunn’s test). Histograms are represented with the means +/-SEM of at least 3 independent experiments. **Figure 5—Figure supplement 3-source data 1** Western blot scans for the data presented in Figure S3b (KIF1A levels in cortical neurons). Shown in red are the cropped regions presented in Figure 5d. Films containing the samples are shown.

**Figure S4.**
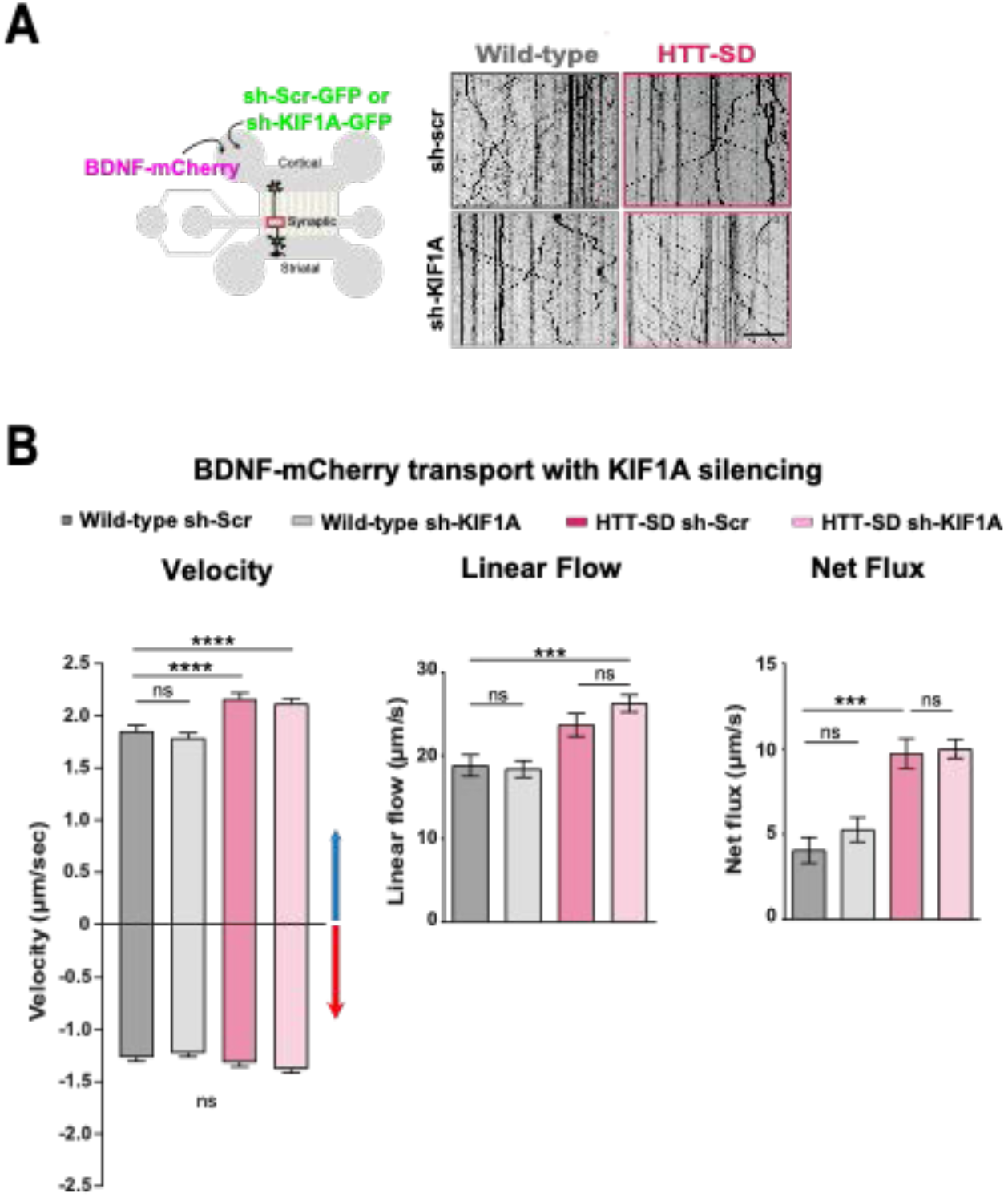
KIF1A silencing doesn’t affect BDNF-mCherry transport. (**A**) Schematic of the 3-compartment microfluidic device with an indication of lentiviral transduction of BDNF-mCherry and sh-scramble (sh-scr-GFP) or sh-KIF1A (sh-KIF1A-GFP) lentiviruses (*left*). Representative kymographs of BDNF-mCherry vesicle transport within WT or HTT-SD axons transduced with sh-Scr or sh-KIF1A (*right*) at DIV8. Scale bar = 25 µm. (**B**) Segmental anterograde (****p<0.0001; N=WT sh-Scr: 618 vesicles, WT sh-KIF1A: 901 vesicles, HTT-SD sh-Scr:1735 vesicles, HTT-SD sh-KIF1A: 2830 vesicles) and retrograde velocities. There were no significant differences between genotypes in the retrograde segmental velocities and KIF1A silencing conditions (ns=non significant), linear flow (*** p<0.001; N=WT sh-Scr: 75 axons, WT sh-KIF1A: 102 axons, HTT-SD sh-Scr: 114 axons and HTT-SD sh-KIF1A: 191 axons), or net flux (*** p<0.001; N=WT sh-Scr: 75 axons, WT sh-KIF1A: 103 axons, HTT-SD sh-Scr: 123 axons and HTT-SD sh-KIF1A: 186 axons) of BDNF-mCherry vesicles. Histograms represent means +/-SEM of 3 independent experiments. Significance was determined using the Kruskal-Wallis test followed by a Dunn’s post-hoc analysis.

